# Catalytic pocket of Clr4 (Suv39h) methyltransferase serves as a substrate receptor for Cullin 4-dependent histone H3 ubiquitination

**DOI:** 10.1101/2025.08.28.672867

**Authors:** Katarina Psenakova, Swapnil S. Parhad, Joao A. Paulo, Xinyue Liu, Emily F. Patterson, Rachel Watson, Marcus A. Cheek, Michael-Christopher Keogh, Marian Kalocsay, Steven P. Gygi, Lucas Farnung, Danesh Moazed

**Affiliations:** Howard Hughes Medical Institute, Department of Cell Biology, Harvard Medical School, Boston, MA, 02115, USA; Institute of Organic Chemistry and Biochemistry of the Czech Academy of Sciences, Prague, Czech Republic; Department of Cell Biology, Harvard Medical School, Boston, MA, 02115, USA; EpiCypher Inc., Durham NC 27709, USA; Department of Experimental Radiation Oncology, University of Texas MD Anderson Cancer Center, Houston, TX 77030, USA

**Author notes:** Correspondence should be addressed to D.M.

## Abstract

Histone H3 lysine 9 (H3K9) methylation must be regulated to prevent inappropriate heterochromatin formation. Regulation of the conserved fission yeast H3K9 methyltransferase Clr4 (Suv39h) involves an automethylation-induced conformational switch and interaction of its catalytic SET domain with mono-ubiquitinated histone H3 lysine 14 (H3K14ub), a modification catalyzed by the Cul4 subunit of the CLRC complex. Using reconstituted CLRC, we show that Clr4 catalytic pocket serves as a substrate receptor for Cul4-dependent H3K14 ubiquitination. H3K14ub activates Clr4 to catalyze cis methylation of H3K9 on the same histone tail, while Clr4 auto-methylation enables H3K14ub-bound Clr4 to methylate H3K9 on an unmodified H3 tail in trans. Crosslinking and structural modeling reveal interactions between Clr4 chromo and SET domains, and between the chromo-domain and H3K14ub, suggesting that the chromodomain reads H3K9me3 and H3K14ub to allosterically regulate Clr4 activity. H3K14 ubiquitination therefore regulates Clr4 by promoting its recruitment and by positioning H3K9 in the active site.

## Introduction

Eukaryotic genomes contain extensive domains of heterochromatin that are required for silencing of transposons and regulation of cell type-specific gene expression ^1,2^. Histone 3 lysine 9 (H3K9) methylation is a conserved feature of heterochromatin which is catalyzed by the Suv39h family of enzymes in organisms from fission yeast to human. The fission yeast Suv39h, Clr4, is a component of a Cullin 4 (Cul4) E3 ubiquitin ligase complex called CLRC, which mono-ubiquitinates H3 lysine 14 (H3K14ub1) ^3–6^. Clr4 contains a ubiquitin binding site adjacent to its catalytic pocket and ubiquitination of K14 appears to activate Clr4 methyltransferase activity by positioning K9 in the catalytic pocket ^7^. The activity of Clr4 is additionally regulated by autoinhibition and its reversal by an automethylation-induced conformational switch that releases a pseudo-substrate lysine from the catalytic pocket of the enzyme to allow histone H3 tail binding and K9 methylation ^8^. The deposition of H3K9 methylation can initiate heterochromatin formation and gene silencing. The activity of H3K9 methyltransferase enzymes must therefore be tightly regulated to prevent inappropriate gene silencing. Indeed, loss of Clr4 autoinhibition leads to spurious heterochromatin formation and deleterious gene silencing ^8^. How the above mechanisms work together to regulate Clr4 activity and heterochromatin formation is not understood.

CLRC shares architectural similarity with other Cullin-RING E3 ubiquitin ligase (CRL) complexes. In addition to Clr4 and Cul4, CLRC contains the Rik1, Raf1, and Raf2 sub-units ^3–5,9^. Rik1 and Raf1 are homologs of DDB1 and DDB2, respectively, which are subunits of canonical Cul4 complexes involved in DNA damage repair ^10^, and like Clr4, Raf2 is a CLRC-specific subunit. In CRL4 complexes, DDB1 acts an adaptor that binds to several substrate receptors including DDB2. By analogy, Rik1 and Raf1 might be expected to act as the adaptor and substrate receptor modules in the CLRC complex, respectively, that promotes H3K14 ubiquitination. Notably, Cul4 has also been reported to ubiquitinate Clr4 to regulate heterochromatin formation ^11^. However, the requirements for specific ubiquitination of H3K14 and Clr4, and the relative contributions of each ubiquitination event to silencing, are not understood. In particular, how Cul4 ubiquitinates a single lysine, K14, in the flexible N terminus of H3 remains unclear.

The coupling of H3K9 methylation and ubiquitination of adjacent lysines in histone H3 appears to be conserved. In mammalian cells, the ubiquitination of H3K18 and K23 mediated by UHRF1, an E3 ubiquitin ligase that binds to CpG dinucleotides and H3K9me3 helps recruit the DNA methyl-transferase DNMT1 and is required for maintenance of DNA methylation ^12,13^. More recently, the ubiquitination of H3K14 by G2E3 ubiquitin ligase has been shown to promote SUV39H recruitment and H3K9 methylation in human cells ^14^. Together these studies suggest a conserved and fundamental role for H3 ubiquitination in regulation of H3K9 methylation and heterochromatin.

In this study, we set out to understand how the CLRC complex mediates the specific ubiquitination of H3K14 and the extent to which H3 and Clr4 ubiquitination contribute to H3K9 methylation and silencing. We expressed subunits and the CLRC complex in insect cells and examined the ubiquitination activity the purified complex on nucleosome substrates. Our results show that the catalytic pocket of Clr4 acts as a substrate receptor for the specific ubiquitination of H3K14, which activates Clr4 for cis methylation of H3K9 on the same histone tail. Strikingly, Clr4 automethylation enables it to methylate H3K9 on an unmodified H3 tail in trans. Cul4 also ubiquitinates Clr4 on multiple lysines and promotes its dissociation from the CLRC complex, which plays a role in spreading of heterochromatin away from nucleation sites. Finally, using crosslinking and mass spectrometry (XL-MS) coupled to structural modeling and other approaches, we identify interactions between the Clr4 chromo and SET domains, and between the chromodomain and H3K14ub. Our findings suggest distinct roles for ubiquitination in regulation of Clr4 activity and heterochromatin formation involving contributions to the initial recognition of H3K9me and H3K14ub1 by the Clr4 chromodomain, the activation of Clr4 methyltransferase activity for cis methylation of H3K9 via interactions with the SET domain, and the release of ubiquitinated Clr4 from the CLRC complex.

## Results

### CLRC complex reconstitution

It has previously been shown that the CLRC complex ubiquitinates H3K14 and this ubiquitination increases the enzymatic activity of Clr4 ^6,7^. However, how the Cul4 E3 ubiquitin ligase subunit selectively ubiquitinates H3K14 remains unknown. To gain insight into the regulation of H3K9 methylation (H3K9me) and H3K14 ubiquitination (H3K14ub), we set out to reconstitute these activities using purified CLRC complex and nucleosomes in vitro. For in vitro reconstitution of the CLRC complex (Fig 1A), we used a baculovirus-insect cell expression system coupled with ligation independent cloning to integrate multiple genes flanked by the PolH promoter and SV40 terminator into the AcMNPV genome ^15,16^. We designed a single bacmid DNA containing 6×His-MBP N-terminally tagged Raf1 and untagged subunits Raf2, Rik1 and Cul4 with each gene under the control of the PolH promoter and SV40 terminator (Fig 1A, Fig S1A). After optimizing expression and purification conditions, the two-step affinity purification yielded a complex of 6×HisMBP-Raf1, Raf2, Rik1 and Cul4. All 4 subunits migrated in a single peak in a size-exclusion chromatography (SEC) column (Fig 1B; Fig S1B, C). We utilized a bacterial expression system to express and purify Clr4 with an N-terminal GST tag (Fig 1A, B). Following purification, we cleaved the GST tag and added approximately a 5-fold molar excess of Clr4 to the four-subunit CLRC complex. SEC was then employed to separate the fully assembled CLRC complex from the excess Clr4 (Fig 1B; Fig S1B, C). Mass spectrometry (MS) analysis of the purified CLRC complex verified the presence of all 5 subunits (Fig S1D).

**Figure 1.**
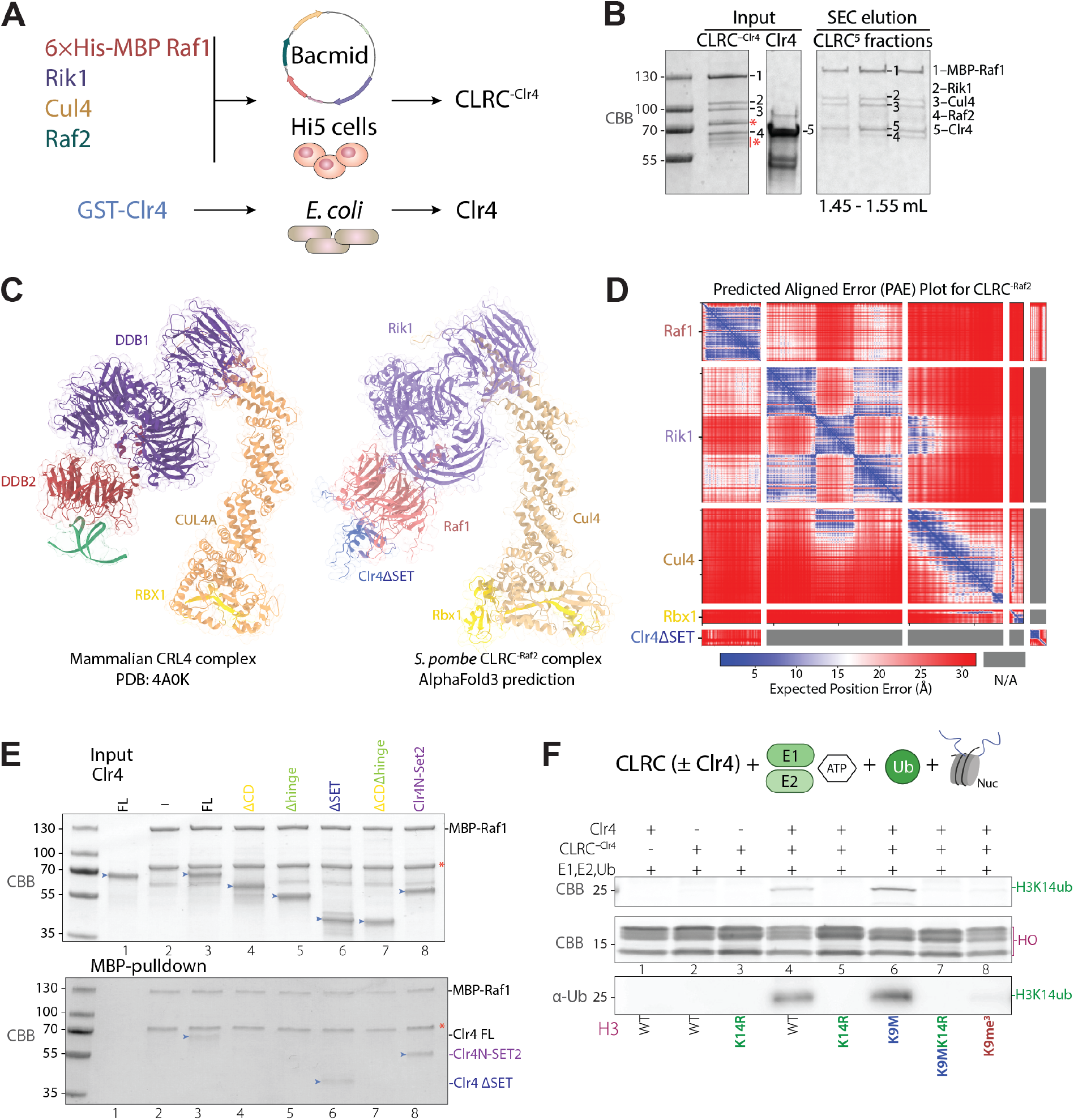
Reconstitution of CLRC complex-mediated H3K14 ubiquitination. **(A)** Strategy for expression and purification of the CLRC E3 complex. The *S. pombe* CLRC complex consists of Cullin 4 (Cul4), the central E3 ligase, Rik1 (DDB1 homolog), Raf1 (DDB2 homolog, Raf2, and the histone methyltransferase Clr4. **(B)** Purification of the five-subunit CLRC complex. SDS-PAGE showing the Ni-NTA purified CLRC complex lacking Clr4 (CLRC^-Clr4^)(left), bacterially expressed and purified Clr4 (middle), and isolation of holo CLRC by size-exclusion chromatography (SEC) on a Superose 6 Increase 3.2/300 column (right). Elution fractions between 1.45-1.55 mL contained all five subunits. *, MBP-Raf1 degradation products. **(C)** Comparison of AlphaFold-predicted structure of CLRC^-Raf2^ with the crystal structure of mammalian CRL4 complex. Crystal structure of the mammalian CRL4 complex (PDB 4A0K) (left); AlphaFold3-predicted structure of the *S. pombe* CLRC^-Raf2^ complex (right). In a pairwise search for interactions, AF3 predicted an interaction between the N-terminal region of Clr4 and Raf1. Protein components are color-coded based on sequence conservation and homology. **(D)** Predicted Aligned Error (PAE) Plot of the AlphaFold3 model of the CLRC complex. PAE plot of the AlphaFold3 model showing predicted alignment confidence between residue pairs, where blue indicates low expected positional error (high confidence), white indicates high expected error (low confidence), and red denotes no predicted interaction between regions. **(E)** Identification of Clr4 domains required for incorporation into CLRC. MBP pulldown assays of CLRC subunits with Clr4 full length (FL) or truncation mutants (light blue arrows). MBP-Raf1 was immobilized on amylose beads and incubated with the indicated Clr4 fragments. After washing and elution, bound proteins were analyzed by SDSPAGE to identify Clr4 domains that are necessary for its binding to MBP-Raf1. *, MBP-Raf1 degradation products. **(F)** Specific ubiquitination of H3K14 by CLRC requires Clr4. SDS-PAGE and Western blot analysis of in vitro ubiquitination assays using nucleosome substrates with the indicated histone mutations. Full ubiquitination reactions contained CLRC, E1 (UBE1), E2 (UbcH5c/UBE2D3), ubiquitin (Ub), and ATP. No H3 ubiquitination was observed in the absence of CLRC (lane 1) or without Clr4 (lanes 2-3). Mono-ubiquitination (~25 kDa band) was detected with WT (lane 4) and H3K9M (lane 6) nucleosomes, but not with H3K14R (lanes 3, 5, 7) or H3K9me3 nucleosomes (lane 8). HO, histone octamer.

To gain insight into the architecture and functional conservation of the *S. pombe* CLRC complex, we used AlphaFold3 (AF3) to predict the structure of a core assembly comprising Raf1, Rik1, Cul4, and a fragment of Clr4 lacking its catalytic SET domain (Clr4ΔSET). This model, referred to as CLRC^-Raf2^ (lacking the accessory subunit Raf2), was then structurally aligned with the crystal structure of the mammalian CRL4-DDB2 complex (Fig 1C, D). The comparison revealed overall structural similarity between the two complexes, further supporting the evolutionary conservation of the CRL4 family. In the AF3 model, Rik1 structurally aligns with DDB1, while Raf1 aligns to DDB2, which is known to serve as the substrate receptor in the mammalian complex ^17^. In a pairwise search for interactions between Clr4 and other CLRC complex subunits, AF3 predicted an interaction between the N-terminal region of Clr4 and Raf1 (Fig 1C, D), suggesting a mechanism of substrate recruitment. To validate the AF3-predicted interaction model between Raf1 and the N-terminal region of Clr4, we generated a series of Clr4 truncations (Fig S1E), including one in which the Clr4 SET domain was replaced with the Set2 SET domain (Clr4NSet2), and assessed their ability to bind Raf1 via Raf1-MBP pull-down assays. Among these proteins, only those retaining the full N-terminal region – including both the chromodomain and the hinge region – were able to interact with Raf1 (Fig 1E, lanes 3, 6, and 8), consistent with the interaction interface in the predicted AF3 model.

### Clr4 is required for the specific ubiquitination of H3K14 in the nucleosome

To assess the ubiquitination activity of the reconstituted CLRC complex, we used ubiquitination reactions containing CLRC with or without Clr4 (referred to as CLRC minus Clr4, CLRC^-Clr4^), recombinant human E1, E2, HA-Ubiquitin protein (labeled as Ub), and the N-terminal peptide of histone H3 (amino acids 1-20, AnaSpec) as the substrate (Fig S1F). We found that the CLRC complex, with or without Clr4, but not the E1 and E2 enzymes alone, efficiently ubiquitinated the H3 N-terminal peptide (Fig S1F). However, the substitution of H3K14 with cysteine (H3K14C) or trimethylation of lysine 9 (H3K9me3) did not abolish ubiquitination of the peptide, indicating that CLRC lacked specificity for ubiquitination of K14 with the H3 N-terminal peptide as a substrate (Fig S1F).

We next used a mono-nucleosome as a substrate for CLRC-dependent ubiquitination (Fig 1F). In contrast to the results with the H3 N-terminal peptide, we found that the CLRC complex lacking Clr4 did not ubiquitinate nucleosomal H3 or other histones (Fig 1F, lane 2). However, upon addition of Clr4 to form the holo CLRC complex, a ~25 kDa Coomassie-stained band, consistent with the size of monoubiquitinated H3, appeared (Fig 1F, lane 4). This reaction was remarkably rapid, with the ubiquitinated H3 band detectable at 2 minutes after the initiation of the reaction (Fig S2B). This band was absent when we used the H3K14R nucleosome as a substrate (Fig 1F, lane 5), indicating that it corresponded to the specific ubiquitination of H3K14 (H3K14ub). Clr4 is therefore required for the specific ubiquitination of H3K14, suggesting that it acts as a substrate receptor for the Cul4 E3 ubiquitin ligase.

### Clr4 catalytic pocket acts as a substrate receptor for specific H3K14ub

To determine the region of Clr4 that is required for H3K14 ubiquitination, we used our panel of Clr4 truncations (Fig 1E, Fig S1E) and assessed their ability to promote H3K14ub in vitro. Only Clr4 fragments retaining both the hinge region and the SET domain supported efficient H3K14 ubiquitination (Fig 2A, lanes 3 and 5), suggesting that the hinge and SET domains may promote H3K14 ubiquitination via the assembly of Clr4 into CLRC and presentation of H3K14 for Cul4-E2-mediated ubiquitination, respectively.

**Figure 2.**
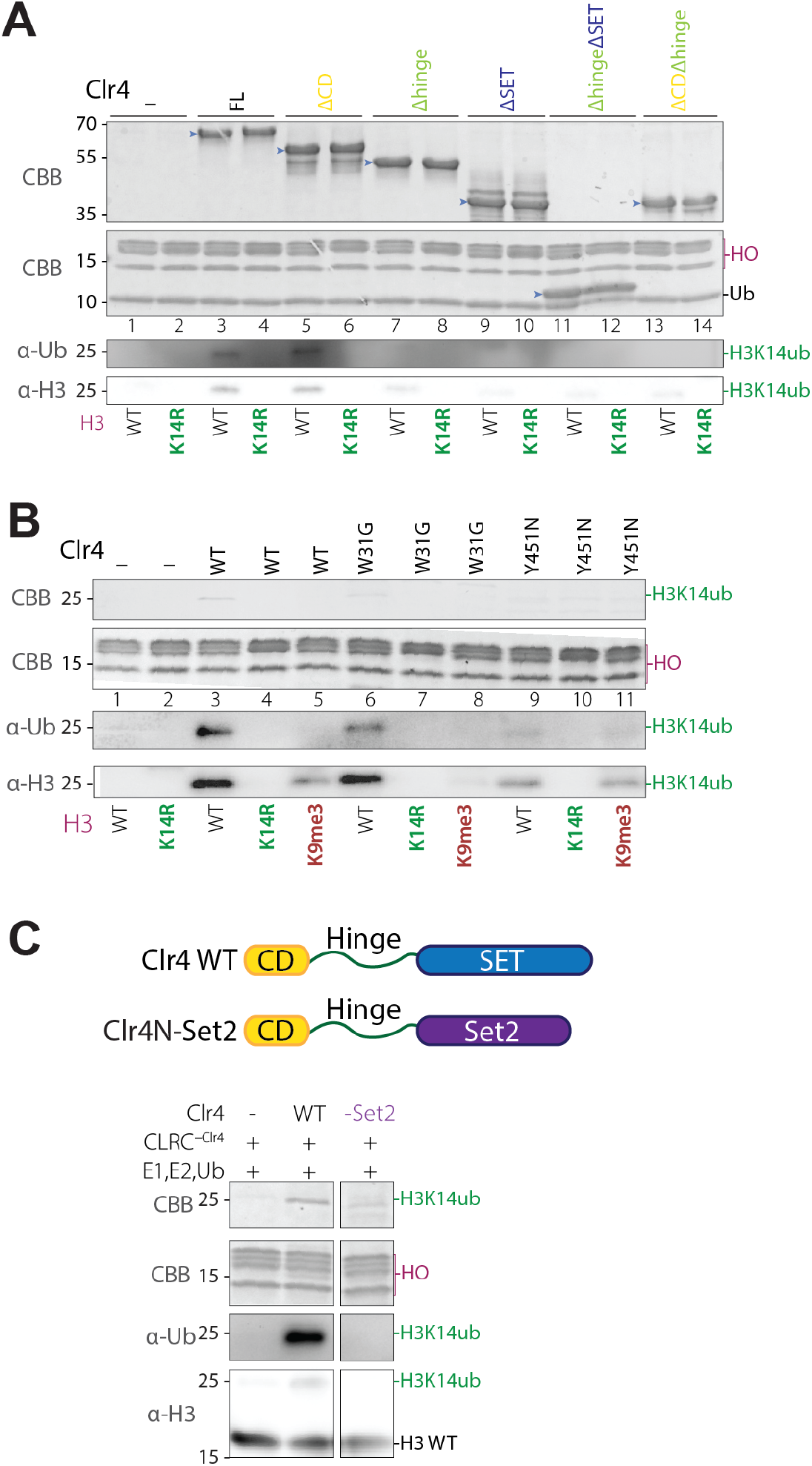
The catalytic pocket of Clr4 acts as a substrate receptor for specific H3K14 ubiquitination. (**A)** Clr4 SET domain is required for H3K14 ubiquitination. In vitro ubiquitination assays with full-length (FL) and the indicated Clr4 fragments (light blue arrows). Only Clr4 fragments retaining both the hinge region and the SET domain allowed H3K14 ubiquitination (lanes 3 and 5). HO, histone octamer. **(B)** Effect of Clr4 mutations and H3K9 methylation on H3K14 ubiquitination. Comparison of WT Clr4 with mutants defective in H3K9me3 binding (W31G, chromodomain) or catalytically dead (Y451N, SET domain). The Y451N mutation markedly impaired H3K14 ubiquitination (lane 9), while W31G caused only a moderate reduction in H3K14 ubiquitination (lane 6). H3K9me3 inhibited H3K14 ubiquitination by both WT and W31G Clr4 proteins. HO, histone octamer. **(C)** Clr4N-Set2 fusion fails to promote H3K14ub. Ubiquitination assays using WT Clr4 or a chimera consisting of the Clr4 chromodomain-hinge fused to the SET domain of *S. pombe* Set2. WT Clr4 promoted H3K14 ubiquitination (lane 2), whereas the Set2 fusion failed to promote ubiquitination (lane 3), suggesting s specific role for the Clr4 SET domain as a substrate receptor for H3K14 ubiquitination. HO, histone octamer.

To examine the relationship between H3K9 methylation and H3K14 ubiquitination, we used a mono-nucleosome substrate with a trimethyled H3K9 (H3K9me3) as a substrate for CLRC-mediated ubiquitination. We found that H3K14 ubiquitination was greatly diminished on H3K9me3 nucleosomes, suggesting that CLRC ubiquitinates H3K14 prior to H3K9 methylation (Fig 1F, lane 8). However, the substitution of H3K9 with methionine (H3K9M), previously shown to lock the catalytic SET domain onto the H3 tail with high affinity ^18^, resulted in more efficient H3 ubiquitination (Fig 1F, lane 6). This H3 ubiquitination was abolished when we used H3K9MK14R nucleosomes as substrate (Fig 1F, lane 7), indicating that H3K9M specifically increased H3K14 ubiquitination. These results demonstrate that the active site of the Clr4 methyltransferase can act as a substrate receptor that presents the H3 tail for specific ubiquitination.

To further examine the relationship between the active site of Clr4 and H3K14 ubiquitination, we reconstituted the CLRC complex using a previously described catalytically inactive Clr4 with a substitution of tyrosine 451 to asparagine (Clr4 Y451N) ^8^. This mutation significantly reduced H3K14 ubiquitination (Fig 2B, lane 9), suggesting that binding and recognition of the H3 N-terminal tail by the catalytic pocket in the SET domain is necessary for efficient H3K14 ubiquitination.

To determine the possible role of Clr4 chromodomain, which binds to the H3K9me3-modified nucleosome, in H3K14 ubiquitination, we reconstituted the CLRC complex containing a Clr4 chromodomain mutant, W31G, which is deficient in binding the H3K9me3-modified histone tail ^19,20^. We found that the Clr4 W31G mutant caused only a moderate decrease in H3K14 ubiquitination (Fig 2B, lane 6) and did not restore H3K14 ubiquitination on an H3K9me3 nucleosome as a substrate (Fig 2B, lane 8). These results suggest that the Clr4 chromodomain does not contribute to H3K14 ubiquitination. Furthermore, the inability of CLRC to ubiquitinate H3K9me3 nucleosomes is not due to the binding of the chromodomain to the H3K9me3 tail, but rather to the inability of the SET domain to bind and recognize an already methylated nucleosome.

As a further test of the requirement for the Clr4 SET domain in specific H3K14 ubiquitination, we designed a fusion construct combining the N-terminal chromodomain and hinge regions of Clr4 with the SET domain of the H3K36 methyltransferase Set2 (Clr4N-Set2; Fig 2C, top). We confirmed that the Clr4N-Set2 fusion protein specifically methylated H3K36, indicating that Set2 retained its substrate methylation specificity in the fusion protein (Fig S2B). Clr4N-Set2 was also incorporated into the CLRC complex (CLRC-Set2, Fig 1E, lane 8). However, unlike CLRC, CLRC-Set2, did not promote nucleosome ubiquitination (Fig 2C), indicating that the specific mode of H3 amino terminus-Clr4 SET domain association is required for ubiquitination of H3K14.

### Clr4 autoinhibition prevents H3K9 methylation of a non-ubiquitinated H3 tail in trans

We previously identified an internal autoregulatory loop (ARL, amino acids 453-472) in Clr4 that inhibits its catalytic activity by obstructing the H3K9 substrate-binding pocket ^8^. This loop undergoes automethylation on specific lysine residues, which triggers a conformational switch that enhances the H3K9 methylation activity of Clr4. Since H3K14 ubiquitination also increases the enzymatic activity of Clr4 ^7^, we tested whether the two mechanisms work together to activate Clr4. We performed methyltransferase assays using both wild-type Clr4 and an automethylation-deficient mutant (Clr4 K455,472R), in which two lysines within the ARL are substituted with arginines to prevent automethylation at these sites, together with reconstituted mono-nucleosomes, including unmodified and H3K14ub-modified nucleosomes, as substrates. Consistent with previous findings ^8^, histone methyltransferase assays using radiolabeled [^3^H]-SAM revealed that the automethylation-deficient Clr4 mutant exhibited reduced H3 methylation activity compared to wild-type Clr4 on unmodified nucleosome substrates (Fig. 3A, lanes 2–3). However, wild-type and automethylation-deficient Clr4 exhibited comparable H3 methylation activity with H3K14ub nucleosomes (Fig 3A, lanes 4-7). In addition, Clr4 showed greatly increased H3 methylation activity on H3K14ub relative to unmodified nucleosomes (Fig 3A, compare lanes 2-3 with 4-7; note longer exposure time for lanes 2-3). These findings indicate that under our reaction conditions, H3K14ub bypasses the requirement for automethylation.

**Figure 3.**
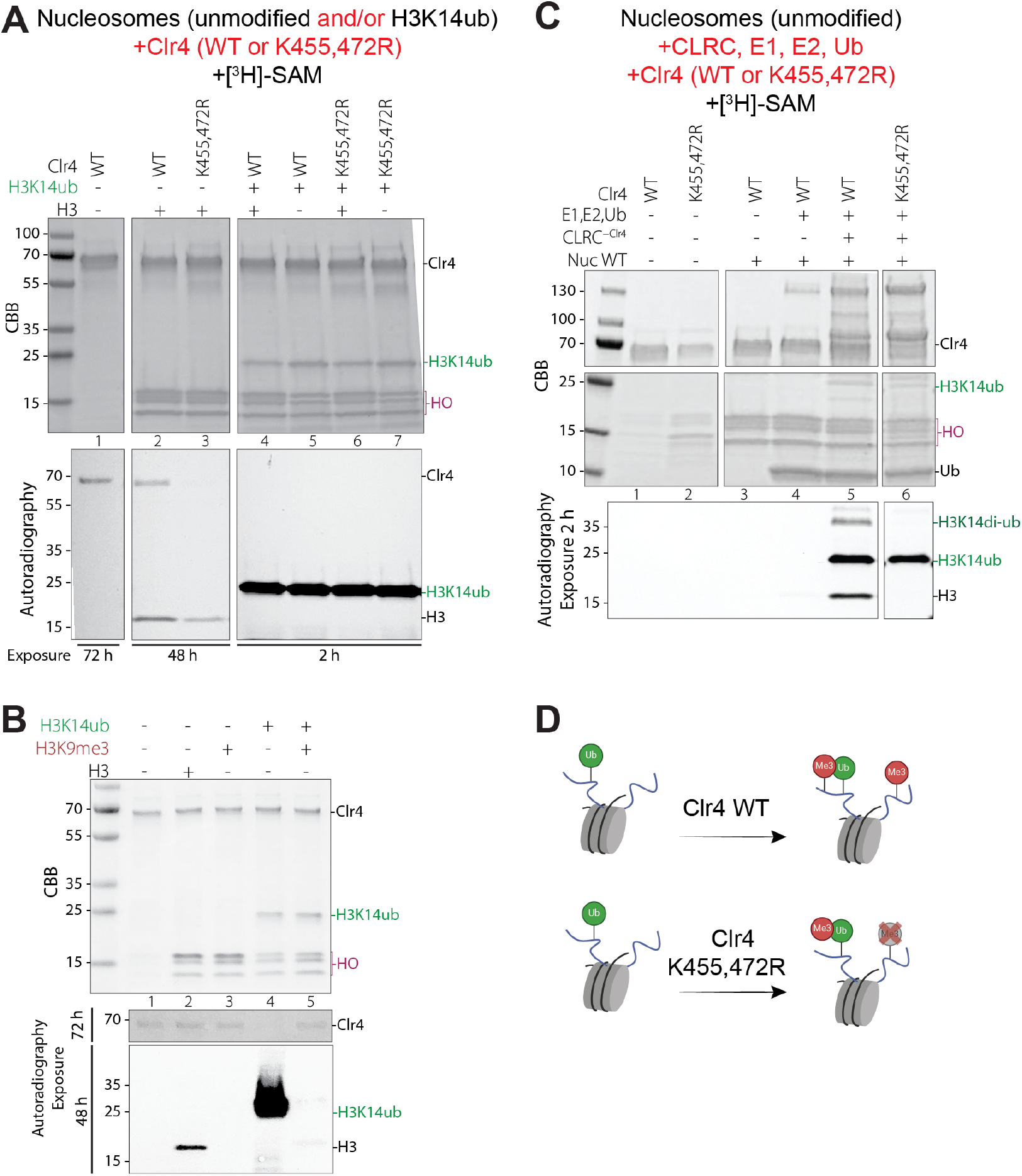
The relationship between Clr4 automethylation and H3K14 ubiquitination. **(A)** H3K14ub activates Clr4 for intranucleosomal methylation independently of its automethylation. Methyltransferase assays with wild-type (WT) and an automethylation-deficient mutant (K455,472R) Clr4 proteins were performed using unmodified and/or H3K14ub-modified nucleosomes. The automethylation mutant showed reduced activity on an unmodified nucleosome (lane 3), but both enzymes displayed similar activity on H3K14ub substrates (lanes 4–7). H3K14ub markedly enhanced Clr4 activity compared to unmodified substrates, despite longer exposure for lanes 2-3. No methylation of unmodified H3 was observed in reactions containing both unmodified and H3K14ub nucleosomes (lanes 4, 6), indicating that under these reaction conditions H3K14ub stimulates intranucleosomal cis H3K9 methylation. HO, histone octamer. **(B)** Clr4 in vitro methyltransferase assay using radioactively labeled [^3^H]-SAM. Methyltransferase assays using Clr4 WT and nucleosomes modified with H3K9me3 or double-modified with H3K9me3 and H3K14ub. No H3 methylation signal was observed with H3K9me3 or doubly modified H3K9me3K14ub nucleosomes (lanes 3 and 5), but Clr4 was automethylated in the doubly modified nucleosome (lane 5). HO, histone octamer. **(C)** H3K14ub promotes Clr4-mediated methylation of K9 on an unmodified H3 tail in an automethylationdependent manner. In vitro methylation and ubiquitination assays were reconstituted using Clr4 WT and Clr4 K455,472R proteins. Both WT and K455,472R Clr4 proteins ubiquitinated H3 (~25 kDa band detected by Coomassie staining, middle panel, lanes 5 and 6). Both WT Clr4 and Clr4 K455, 472R efficiently methylated a ubiquitinated form of H3, but only WT Clr4 methylated unmodified H3 (autoradiography, lower panel, lanes 5 and 6). HO, histone octamer. **(D)** Schematic summary based on the results in panels A-C.

A recent cryo-EM structure of the Clr4 SET domain bound to an H3K14ub-modified N-terminal histone peptide ^21^ demonstrates that the binding of the Clr4 SET domain to ubiquitin releases the ARL. This is consistent with our observation that H3K14ub can bypass the requirement for Clr4 automethylation on a nucleosomal substrate (Fig 3A, lane 7). However, in our reconstituted reactions, we observed Clr4 automethylation in the presence of H3K9me3K14ub nucleosomes (Fig 3B, lane 5). The ubiquitin-mediated release of the ARL in the SET domain may therefore be transient, potentially allowing automethylation to occur.

To gain insight into the interplay between autoregulation and H3K14ub-dependent regulation of Clr4 activity, we established a fully reconstituted *in vitro* system that included both enzymatic reactions. Clr4 alone, in the absence of the CRL scaffold, did not methylate unmodified nucleosomes (Fig 3C, lanes 1-4). Surprisingly, in the fully reconstituted system using Clr4-CLRC, we observed methylation of both H3K14ub-modified and unmodified nucleosomes, suggesting that Clr4 bound to a ubiquitinated nucleosome could methylate an H3 tail that had not yet been ubiquitinated (Fig 3C, lane 5). However, when we performed the ubiquitination and methylation reactions using Clr4 K455,472R, which cannot be automethylated, CLRC-Clr4-K455,472R efficiently methylated the H3K14ub-modified nucleosome substrate but not nucleosomes containing non-ubiquitinated H3 (Fig 3C, lane 6), suggesting that Clr4 autoinhibition restricts H3K9 methylation on non-ubiquitinated H3 tails in trans. To test whether this trans-methylation occurs intraor inter-nucleosomally, we performed reactions containing an equimolar mixture of fully unmodified and fully H3K14 ubiquitinated nucleosomes (Fig 3A, lanes 4 and 6). As shown in Fig 3A, lane 4, H3 was methylated only in the H3K14ub but not the unmodified nucleosomes in the mixture, indicating that the trans H3K9 methylation occurs intra-nucleosomally. Taken together, these findings suggest that automethylation allows Clr4 to act in trans intra-nucleosomally, engaging one H3 tail when the other is ubiquitinated (Fig 3D).

### Regulation of Clr4 activity through a second ubiquitin binding site

We were interested in understanding whether the interaction of the SET domain of Clr4 with H3K14ub was sufficient for binding of Clr4 to H3K14ub nucleosomes. We tested the ability of full length Clr4, Clr4Δhinge (lacking the hinge domain), and Clr4-SET (ΔCDΔhinge, lacking the chromodomain and hinge domains) to form complexes with H3K14ub nucleosomes. The results showed that full-length Clr4 and Clr4Δhinge co-migrated with H3K14ub nucleosomes during size exclusion chromatography (SEC), suggesting that they formed stable complexes (Fig 4A). By contrast, Clr4-SET did not co-migrate with H3K14ub nucleosomes, suggesting that interactions involving the Clr4 chromodomain with the ubiquitinated nucleosome are required to stabilize the complex.

**Figure 4.**
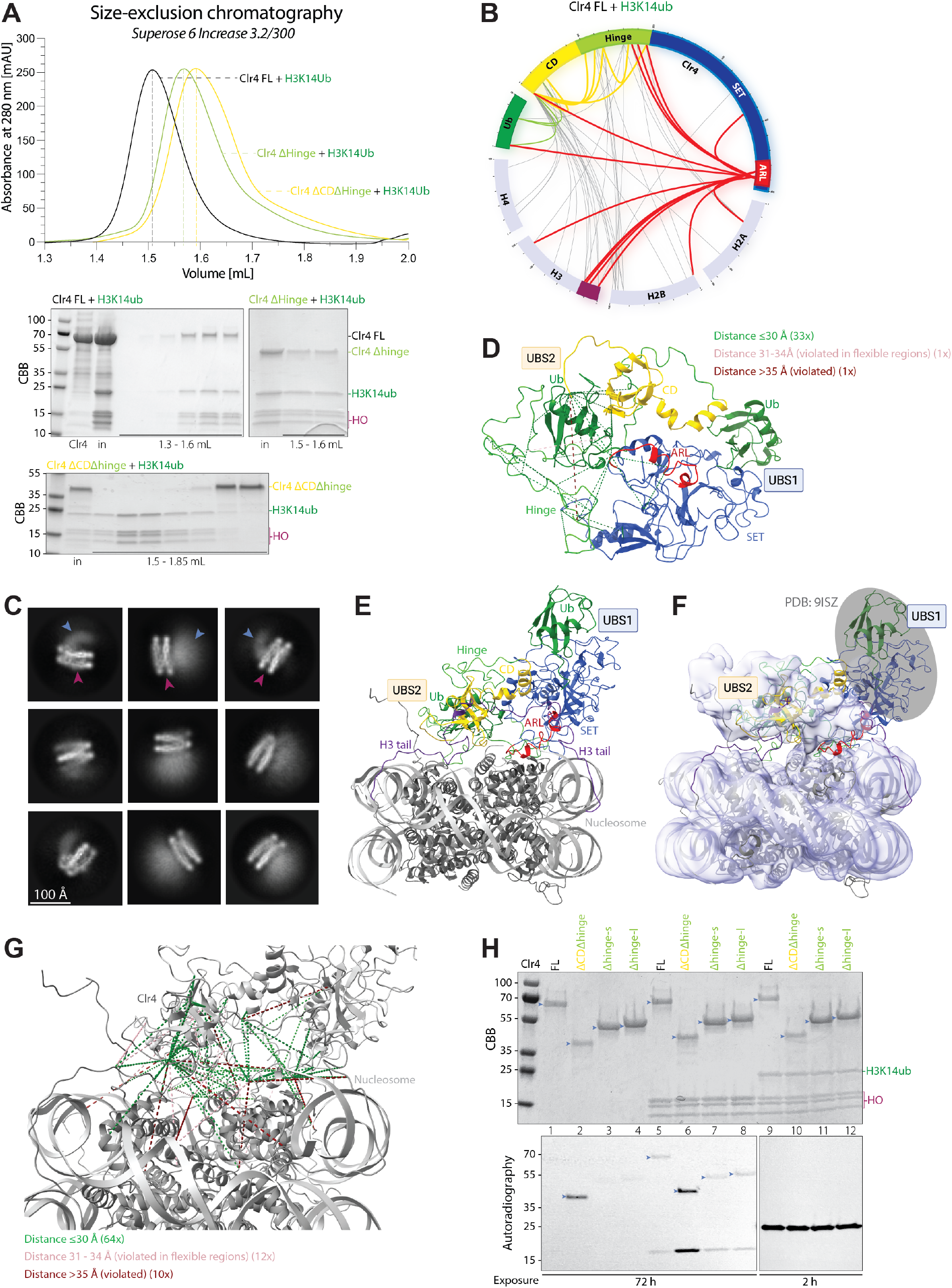
Structural and functional basis of Clr4 recognition of H3K14ub-modified nucleosomes. **(A)** The chromodomain promotes Clr4 binding to H3K14ub nucleosomes. Size-exclusion chromatography (SEC) analysis of complex formation between the indicated full-length (FL) Clr4 or its subfragments and H3K14ub-modified nucleosomes. Full-length Clr4 and a hinge domain-deleted variant (Clr4Δhinge) co-migrated with H3K14ub nucleosomes, suggesting the formation of stable complexes. By contrast, the Clr4-SET, which lacks both the chromodomain and hinge region (ΔCDΔhinge), did not co-migrate with H3K14ub nucleosomes, suggesting that the chromodomain is essential for stable nucleosome binding in the context of H3K14 ubiquitination. HO, histone octamer. **(B)** Cross-linking mass spectrometry reveals a shared interface between ubiquitin, Clr4 chromodomain, and the autoregulatory loop (ARL). Cross-linking mass spectrometry (XL-MS) was performed on the full-length Clr4-H3K14ub nucleosome complex. Cross-linked lysine residue pairs were mapped onto the linear domain architecture of Clr4, histones, and ubiquitin, and visualized as connecting lines. Intermolecular cross-links between the Clr4 chromodomain and ubiquitin (green lines) or Clr4 autoregulatory loop (ARL, red), and between the chromodomain, hinge region, and ubiquitin (red lines) are highlighted. The histone H3 tail (highlighted in violet) displayed numerous cross-links with both Clr4 chromodomain and ARL. For clarity, only crosslinks involving Clr4 are shown; all crosslinks, including those within the nucleosome, are listed in Table S5. **(C)** Representative 2D class averages show extranucleosomal density on top of the nucleosomal disk (light blue arrows, scale bar 100 Å). An extended set of 2D classes is shown in Fig S3B. Nucleosome highlighted by dark-pink arrows. **(D)** XL-MS-guided model of Clr4 in complex with two ubiquitin molecules via ubiquitin interactions site 1 (UBS1, blue) and ubiquitin interaction site 2 (UBS2, yellow). The Clr4ubiquitin complex model was generated using HADDOCK, with input from an AlphaFold3 model of pairwise Clr4-ubiquitin interactions (see Figure S4), guided by crosslinking mass spectrometry (XL-MS) restraints. The model satisfies 33 crosslinks within a 30 Å threshold, with one mild violation (31-34 Å) in a flexible region and one significant violation (>35 Å). **(E)** Model of Clr4-H3K14ub nucleosome interaction guided by XL-MS restraints. HADDOCK docking driven by XL-MS distance constraints was used to model the interaction between Clr4 and the H3K14ub nucleosome, using AF3 and XL-MS guided HADDOC model of Clr4 (panel C) and the nucleosome (PDB 1KX5) as input. The resulting complex was refined by molecular dynamics in YASARA. The model suggests that the Clr4 chromodomain (CD, yellow) and autoregulatory loop (ARL, red) interact directly with the nucleosome core and with H3K14 ubiquitinated H3 tail via UBS1 and UBS2; Ub, green, SET domain, blue. **(F)** CryoEM map of the Clr4-H3K14ub nucleosome complex. The Clr4-H3K14ub nucleosome model was fitted into a low-pass filtered cryo-EM map (purple surface representation, 50% transparency) using COOT (see Figure S4). The density observed at the top of the nucleosome likely corresponds to a composite of the Clr4 chromodomain, one ubiquitin moiety (UBS2), and the ARL region (color coding as in panel E). The dynamic nature of these regions likely contributes to the low-resolution features of the map. The second ubiquitin and the unresolved portions of the SET domain (UBS1, docked from PDB 9ISZ), which could not be confidently fitted due to flexibility, are outlined in gray. **(G)** Mapping of XL-MS crosslinks onto the Clr4-H3K14ub nucleosome model. Of the 86 crosslinks identified in the XL-MS analysis, the majority were satisfied within expected distance thresholds, with distances ≤30 Å highlighted in green and 31-34 Å in pink. Ten crosslinks exceeded 35 Å (dark red). Six of these longer-distance crosslinks involved the flexible N-terminal linker and hinge regions of Clr4, while the remaining four connected flexible histone tails to Clr4. These outliers likely reflect conformational flexibility and dynamic regions not fully represented in the static model. **(H)** Methyltransferase activity of full length and truncated Clr4 proteins (light blue arrows) with unmodified and H3K14ub nucleosomes. In vitro methyltransferase assays performed using the indicated full-length or truncated Clr4 proteins and reconstituted nucleosomes containing either unmodified H3 (WT, lanes 5-8) or ubiquitinated H3K14 (H3K14ub, lanes 9-12). Deletion of the Nterminal chromodomain (CD) enhanced Clr4 automethylation (lanes 2 and 6) and H3K9 methylation (lane 6), suggesting that the chromodomain inhibits automethylation and H3 substrate methylation. HO, histone octamer.

To analyze the interaction between full-length Clr4 and ubiquitinated nucleosomes, we employed cross-linking coupled with mass spectrometry (XL-MS) using the amine-reactive cross-linker bissulfosuccinimidyl suberate (BS3), which links lysine side chains or N-terminal α-amino groups. Notably, the region of the SET domain, previously shown to interact with ubiquitin (here referred to as Ubiquitin-binding site 1, UBS1) ^7,21^ lacks lysine residues, and as expected, we did not detect any cross-links between the folded region of the SET domain and ubiquitin (Fig 4B). Surprisingly, we observed multiple cross-links between the Clr4 chromodomain and ubiquitin (Fig 4B, green) as well as between the Clr4 ARL and the chromodomain, hinge region, and ubiquitin (Fig 4B, red). These results suggest a shared binding interface between ubiquitin, the chromodomain and ARL, and the presence of a second ubiquitin-binding site within Clr4, here referred to as Ubiquitin-binding site 2 (UBS2), in addition to UBS1, the previously identified SET-ubiquitin interface. In addition to the expected cross-links between the Clr4 hinge and SET domains and the H3 N-terminal tail, we observed extensive cross-links between the chromodomain, hinge region, ARL and the nucleosome core. In contrast, no cross-links were detected between the folded regions of the SET domain and the nucleosome core. This suggests that the chromodomain and the ARL are in close proximity to the nucleosome core, but the SET domain appears oriented away from the core except for the ARL.

We next examined Clr4-H3K14ub nucleosome complexes by cryo-EM, but despite extensive efforts, were unable to obtain a high-resolution structure. Our cryo-EM analysis showed that although ~45% of the particles were nucleosomes with clear extra density, the density exhibited too much flexibility and lacked distinct features, which prevented confident assignment of structural elements (Fig S3A-G). However, the data showed that all the collected particles exhibited extra density on only one side of the nucleosome (Fig 4C; light-blue arrows, extra density; dark-pink arrows, nucleosome; extended set shown in Fig S3B). This observation contrasts with the structural features of the Clr4-H3K9me3 nucleosome complex in a recently published cryo-EM model ^22^. This difference suggests that the presence of H3K14ub may alter the mode of binding, enabling a single Clr4 molecule to interact with both H3 tails of a single nucleosome.

To model the interaction between Clr4 and a nucleosome, we implemented restrictions driven by XL-MS and AF3 structural predictions. The AF3-predicted model, based on pairwise interactions between Clr4 and ubiquitin, closely recapitulated the SET-ubiquitin interface UBS1 in the recently solved X-ray structure ^21^ (PDB 9ISZ; Fig S4A-C). While AF3 did not predict a chromodomain-ubiquitin interaction, the predicted model could accommodate the chromodomain-ARL crosslinks observed in XL-MS (Fig S4A, D, E). Notably, the ARL adopted an open conformation in the AF3 model compared to the crystal structure of the SET domain (PDB 6BP4; Fig S4F, G). This initial AF3 model served as the input for HADDOCK modeling, incorporating XL-MS distance restraints, and suggested the location of the second ubiquitin-binding interface guided by these restraints (Fig 4D). We also used AF3 to predict interactions between one Clr4 and two H3 N-terminal tails (Fig S4H, I, J). In the AF3 model, one H3 tail was bound to the chromodomain, closely matching the X-ray structure of the *S. pombe* Chp1 chromodomain-H3 tail and the second H3 tail was bound to the Clr4 SET domain-H3 tail, closely matching the X-ray structure of the complex (Fig S4I, comparison with PDB 3G7L and PDB 9ISZ). Our attempts to model the nucleosome-Clr4 interface using AF3 did not yield reliable results.

To model the interaction between Clr4 and the H3K14ub nucleosome, we performed HADDOCK docking driven by XL-MS distance restraints, using our Clr4 model (Fig 4D) and the nucleosome (PDB 1KX5) as input. The resulting complex was relaxed by molecular dynamics in YASARA and suggests that the Clr4 CD and ARL interact directly with the nucleosome core (Fig 4E). This model was then fitted into the low-pass filtered cryo-EM map using COOT (Fig 4F), suggesting that the additional cryo-EM density observed at the top of the nucleosome likely corresponded to a composite of the chromodomain, ubiquitin from one of the ubiquitinated H3 tails, and the ARL region (Fig 4E, F). These regions are likely to be dynamic leading to the low resolution cryo-EM map. We were unable to confidently fit the remainder of the SET domain or the second ubiquitin into the cryo-EM map, presumably due to its inherent flexibility (Fig 4F, outlined in gray). However, this missing region includes the previously published structure of UBS1 ^21^, which we incorporated into the model (Fig 4E, F, PDB 9ISZ). In the resulting model, out of the 86 cross-links identified in our XL-MS analysis, nine exhibited distances between 36-45 Å, and one extended to 56 Å (Fig 4G). Notably, six of these violated cross-links involved the N-terminal flexible linker and hinge regions of Clr4, while the remaining four violations connected flexible histone tails to Clr4 (Fig 4G, Table S1). All crosslinks, including those within the nucleosome, are listed in Table S5.

The proximity of the chromodomain (CD) and the ARL regions of Clr4 in the Clr4-nucleosome model suggests that the chromodomain may regulate the methyltransferase activity of Clr4. To test this hypothesis, we assessed the methyl-transferase activity of full-length Clr4, Clr4Δhinge (lacking the hinge domain), and Clr4-SET (lacking both the chromo-domain and hinge domains; ΔCDΔhinge) using both wild-type and H3K14ub-modified nucleosomes as substrates. Deletion of the chromodomain (Clr4ΔCD) resulted in a marked increase in both Clr4 automethylation and substrate methylation (Fig 4H, lanes 1, 3, 4 and 5, 7, 8, respectively), suggesting that the chromodomain inhibits Clr4 catalytic activity. However, the presence of H3K14 mono-ubiquitination enhanced substrate methylation to comparable levels by both wild-type and Clr4 variants lacking the chromodomain (Fig 4H, lanes 9-12), effectively suppressing the inhibitory effect of the chromodomain. These findings suggest that the chromodomain directly inhibits Clr4 automethylation and H3 substrate methylation through interactions with the ARL region and that this inhibition can be reversed by H3K14ub.

### Clr4 is ubiquitinated in a Cul4-dependent manner

Autoubiquitination of other CRL components by Cul4 is known to regulate the stability and function of these complexes ^23^. To investigate whether CLRC is subject to a similar regulatory mechanism, we reconstituted an *in vitro* ubiquitination reaction using purified CLRC complex. This assay revealed a distinct slower migrating species during gel electrophoresis, which was recognized by an anti-ubiquitin antibody by western blotting and was consistent with the distinct sizes of mono- or di-ubiquitinated Clr4 (Fig S5A, lane 7). These slower migrating species were dependent on the presence of full CLRC complex, and were absent with the E1 and E2 enzymes alone, or in the CLRC^-Clr4^ reaction (Fig S5A, lanes 5,6), suggesting that Cul4 mediated Clr4 ubiquitination. We also tested whether Clr4 was ubiquitinated in vivo in a Cul4-dependent manner. We transformed *S. pombe* cells with a plasmid expressing a 6×His-tagged ubiquitin (6×His-Ub) and conducted Ni-NTA pulldown under denaturing conditions to isolate ubiquitinated proteins in wild-type (WT) and *cul4-1* (C-terminally GFP-tagged loss of function mutation) cells that also expressed FLAG-Clr4 (Fig 5A, B). The *cul4*^*+*^ gene is essential for viability in fission yeast, but *cul4-1* is a viable hypomorphic mutant, previously shown to have a silencing defect ^3^. Using this approach, we detected ubiquitinated FLAG-Clr4 in WT but not in *cul4-1* mutant cells (Fig 5C), indicating that Clr4 was ubiquitinated in a Cul4-dependent manner *in vivo*.

**Figure 5.**
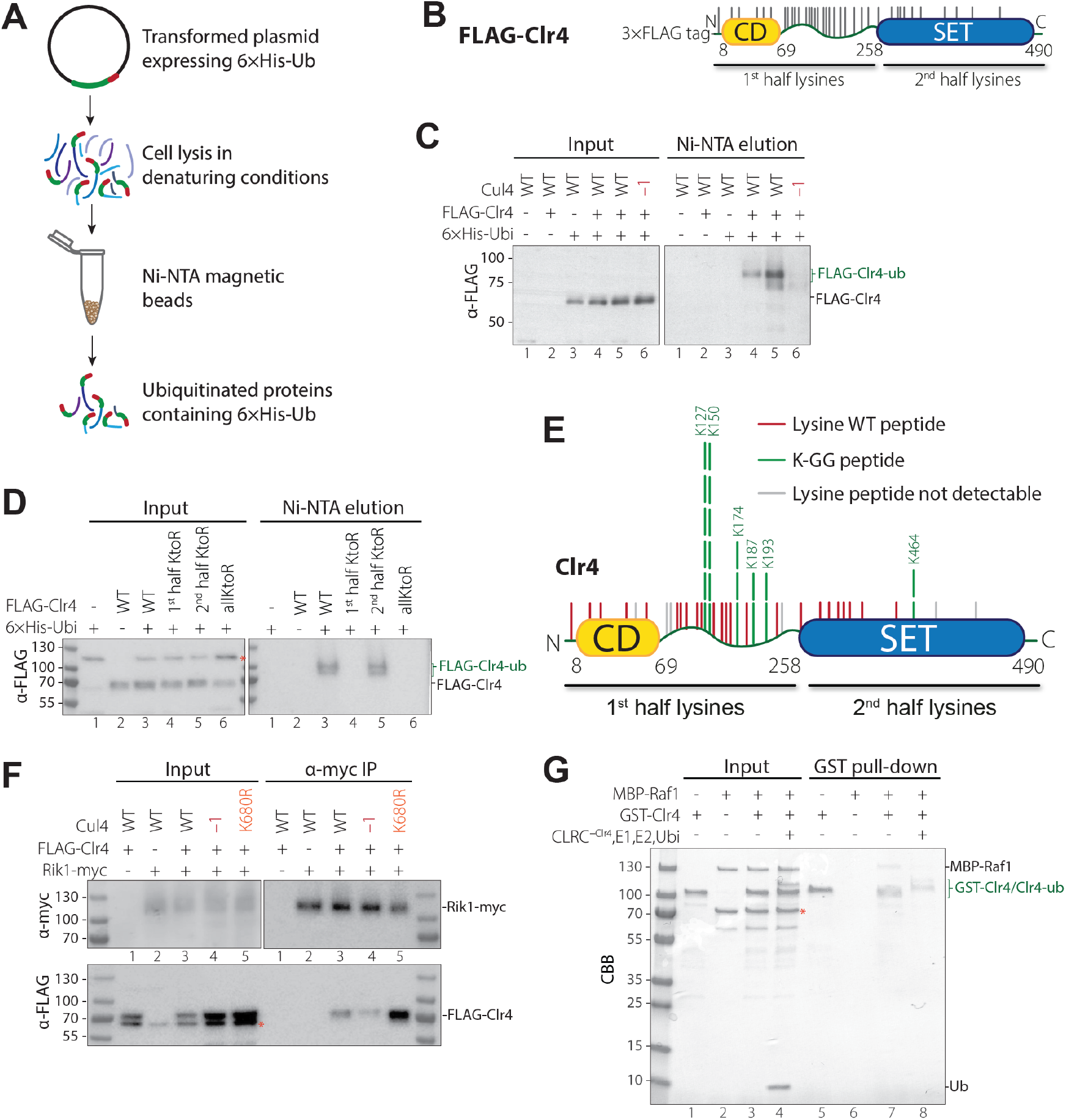
Cul4-mediated Clr4 ubiquitination. **(A)** Overview of the ubiquitinated protein pull-down workflow. Cells were transformed with a plasmid expressing 6×His-tagged ubiquitin (6×His-Ub) and lysed under denaturing conditions to preserve covalent ubiquitin conjugates. Lysates were incubated with Ni-NTA magnetic beads to selectively enrich for ubiquitinated proteins carrying 6×His-Ub. The captured proteins were then subjected to downstream analysis. **(B)** Schematic diagram of the FLAG-Clr4 protein highlighting the location of lysines in the 1^st^ half and 2^nd^ half of the protein used for mutational analysis. Clr4 was expressed with an N-terminal 3×FLAG epitope tag. A total of 36 lysine residues in Clr4 were divided into two groups: 26 lysines located in the N-terminal half, encompassing the CD and hinge region (1^st^ half lysines) and 10 lysines in the C-terminal half, residing within the SET catalytic domain (2^nd^ half lysines). This classification was used to guide targeted mutagenesis and functional assays. **(C)** Clr4 ubiquitination in vivo is dependent on Cul4. Anti-FLAG western blot following Ni-NTA pulldown under denaturing conditions from *S. pombe* cells coexpressing 6×His-tagged ubiquitin (6×His-Ub) and FLAG-Clr4. Ubiquitinated FLAG-Clr4 was detected in wild-type (WT) cells but was greatly diminished in *cul4-1* mutants, indicating that Clr4 ubiquitination requires Cul4 in vivo. **(D)** Clr4 ubiquitination targets lysines in the chromodomain and hinge region. Anti-FLAG western blot following Ni-NTA pulldown under denaturing conditions from *S. pombe* cells co-expressing 6×His-tagged ubiquitin (6×His-Ub) and the indicated FLAG-tagged Clr4 proteins. Cells expressing Clr4 WT or the 2^nd^ half KtoR mutant (lysines mutated in the SET domain) retained ubiquitination. By contrast, Clr4 1^st^ half KtoR (lysines mutated in the chromodomain and hinge region) and Clr4 allKtoR mutants showed loss of ubiquitination, indicating that ubiquitin conjugation occurs primarily within the N-terminal half of Clr4. *, background band present in untagged cells recognized by the anti-FLAG antibody.

Clr4 contains 36 lysines, 26 located in the chromodomain (CD) and the hinge region, and 10 within the SET domain (Fig 5B). To generate Clr4 protein that could not be ubiquitinated on lysine residues, we created a Clr4 mutant with all 36 lysines substituted with arginines (Clr4^allKtoR^). Additionally, we generated two other mutant proteins, Clr4 1^st^ half KtoR and Clr4 2^nd^ half KtoR, in which lysines in the CD and hinge regions or the SET domain were substituted with arginine, respectively (Fig 5B). We transformed cells that express wild-type or the mutated Clr4 proteins with the plasmid expressing 6×His-Ub and performed Ni-NTA pulldown under denaturing conditions. While both wild-type Clr4 (Clr4 WT) and Clr4 2^nd^ half KtoR retained ubiquitination, ubiquitination was lost in Clr4 1^st^ half KtoR and Clr4^allKtoR^ mutants (Fig 5D), indicating that ubiquitination primarily targets the lysines within the CD and hinge region.

We were unable to identify ubiquitinated lysines in Clr4 using native FLAG-Clr4 immunoprecipitation (Fig S5A) and mass spectrometry analysis, likely due to limited amounts of Clr4. We therefore used the *in vitro* reconstitution system with purified CLRC to search for ubiquitinated lysines in Clr4 by mass spectrometry. Following trypsin digestion, ubiquitinated lysines can be identified by their attached GlyGly C-terminal ubiquitin residues. This analysis revealed six ubiquitination sites within Clr4, five of which were localized to the 1^st^ half of Clr4 (Fig 5E), consistent with our *in vivo* ubiquitination results (Fig 5D). The most frequent ubiquitination sites were lysines 127 and 150, both situated within the N-terminal portion of the hinge region. A recent study using Clr4 mobility shift in western blots of cells carrying wild-type Clr4 or Clr4 K to R substitutions reported monoubiquitination of Clr4 on lysines 109, 110, 113, and 114. By contrast, western blot analysis of our FLAG-Clr4 purifications from *S. pombe*, performed in the presence of the proteasome inhibitor MG-132 and the deubiquitinase inhibitor NEM, revealed slower migrating Clr4 species that suggested Clr4 polyubiquitination (Fig S5B). This discrepancy may be due to the different methods used for the identification of ubiquitinated Clr4 species and it also remains possible that Clr4 ubiquitination is not confined to a single lysine and may involve multiple lysines within the flexible hinge region.

Because the Clr4 hinge region is required for its incorporation into the CLRC complex (Fig 1D), we investigated whether Clr4 ubiquitination affects CLRC assembly. The abundance of Clr4 was not affected by mutations that impaired its ubiquitination (Fig 5C, D, F; Fig S5C), but co-immunoprecipitation experiments showed that loss of ubiquitination in *cul4-K680R* mutant cells, in which Cul4 Neddylation and ubiquitination activity are blocked, resulted in increased FLAG-Clr4 association with Rik1-myc (Fig 5F). Consistent with this result, analysis of FLAG-Clr4 immunoprecipitations using quantitative mass spectrometry showed that in *cul4-K680R* mutant cells FLAG-Clr4 associated with about 3-fold more of each of the other CLRC subunits (Fig S5D, green). Interestingly, the hypomorphic *cul4-1* allele, which impairs Cul4-mediated ubiquitination, reduced the interaction of FLAG-Clr4 with Rik1 or its assembly into the CLRC complex (Fig 5F, lane 4; Fig S5D, red). These results suggest that Cul4-mediated Clr4 ubiquitination inhibits its assembly into the CLRC complex. The slight decrease in assembly of Clr4 into CLRC in *cul4-1* cells is likely due to a dominant negative steric inhibition of Clr4 incorporation into CLRC, rather than loss of Clr4 ubiquitination. Consistent with these results, Clr4 1^st^ half KtoR substitutions, which lead to loss of its ubiquitination, led to increased coimmunoprecipitation of FLAG-Clr4 with the Rik1 subunit of CLRC (Fig 5F, Fig S5C). Furthermore, we found that the interaction between Clr4 and Raf1 was abolished following the Cul4-dependent ubiquitination of Clr4 *in vitro* (Fig 5G), suggesting that ubiquitination of Clr4 promotes its release from the CLRC complex.

### Role of ubiquitination in heterochromatic formation

To analyze the relative contributions of Clr4 and H3K14 ubiquitination to heterochromatin formation, we analyzed mutations that selectively disrupt each modification individually, as well as mutations in Cul4, which are expected to eliminate both modifications simultaneously. Specifically, we examined H3K14R and H3K14A substitutions, which block H3K14 ubiquitination, and Clr4 lysine-to-arginine (KtoR) mutants, which impair Clr4 ubiquitination. To evaluate the combined loss of both modifications, we analyzed two *cul4* alleles: the hypomorphic *cul4-1* allele (*cul4-GFP*), which retains partial ubiquitination activity, and the *cul4K680R* allele, which abolishes Cul4 neddylation and substrate ubiquitination. ChIP-seq analysis showed that the association of FLAG-Clr4 with all heterochromatic loci, except the rDNA repeats, was dramatically reduced in *cul4-K680R* cells (Fig 6A). In contrast, *cul4-1* had a more variable effect on FLAG-Clr4 localization, which was increased at the centromeric *dg* and *dh* repeats but reduced or completely lost at the mating type locus and subtelomeric regions (Fig 6A). We observed a weak FLAG-Clr4 enrichment at RNAi nucleation centers in both *cul4* mutant backgrounds (Fig 6A). In *cul4-1* cells, FLAG-Clr4 remained associated with the centromeric *dg/dh* repeats and the mating type *cenH* region, although this association was lower in *cul4-K680R* cells (Fig 6A). These findings suggest that Cul4-mediated ubiquitination is not required for the initial recruitment of Clr4 to nucleation sites but plays a critical role in Clr4 spreading beyond nucleation sites.

**Figure 6.**
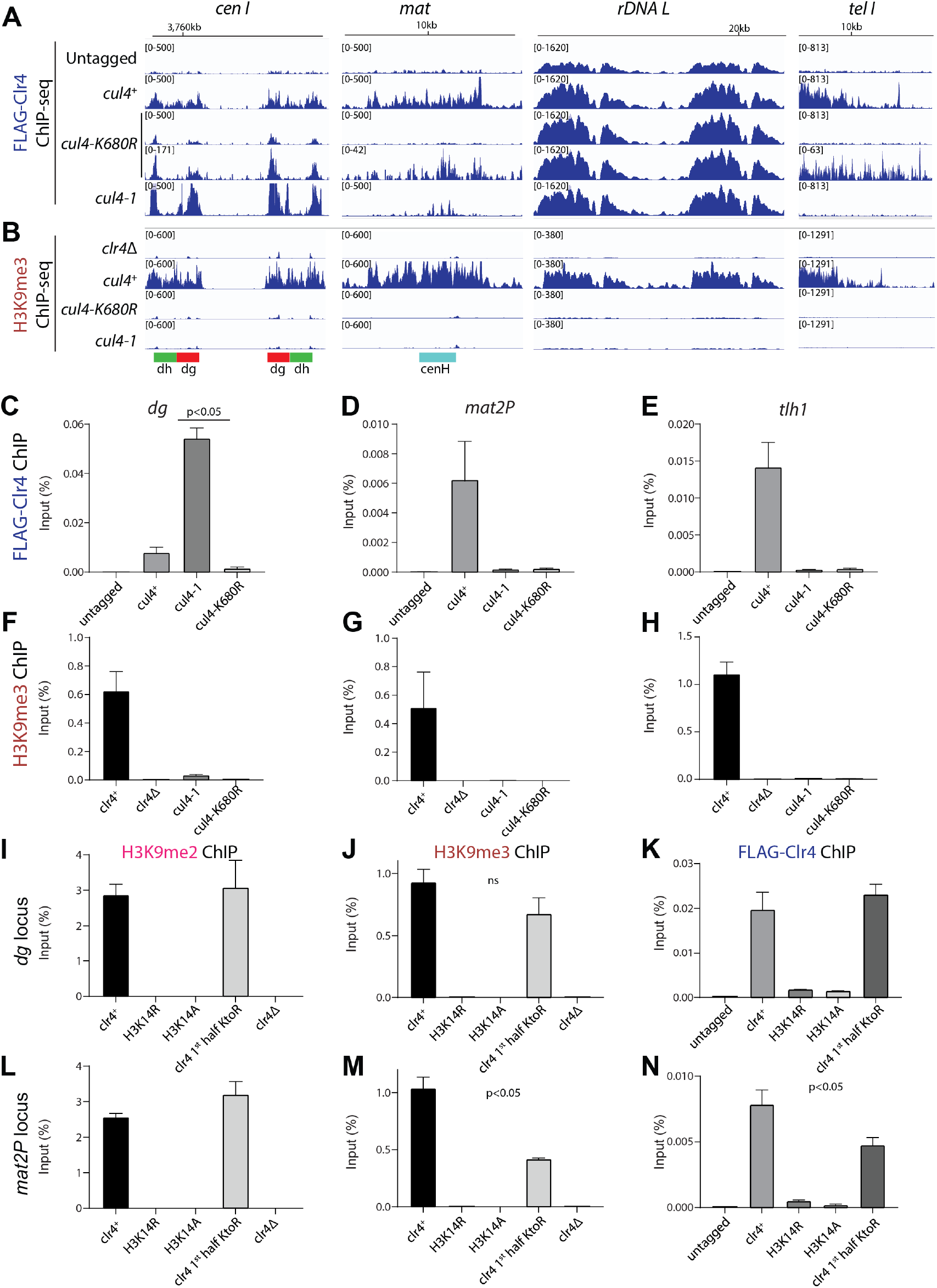
Role of Cul4-mediated ubiquitination in Clr4 localization and H3K9 methylation. **(A, B)** ChIP-seq genome browser tracks showing localization of FLAG-Clr4 (A) and H3K9me3 (B) modification at the indicated (top) genomic loci in wild-type (*cul4*^*+*^), *cul4-K680R*, and *cul4-1* cells. The locations of centromeric *dg*, and *dh* repeats and the mating type (*mat*) centromere homology (*cenH*) are highlighted in the bottom. Chromosome coordinates shown on the top, genotypes on left, and reads per million in brackets. **(C-N)** ChIP-qPCR analysis of FLAG-Clr4, H3K9me2, and H3K9me3 localization at the indicated loci (*dg, mat2P*, and *tlh1*) in wild-type, *cul4-K680R, cul4-1, H3K14R, H3K14A, 1*^*st*^ *half-KtoR* (clr4 with 1^st^ half lysines substituted with arginine, see Figure 5B), and *clr40* cells. Bars show mean percentage input and error bars show standard deviations of 3 biological replicates.

We also examined H3K9me3 in *cul4* mutant cells, using *clr4Δ* as a negative control. The results showed that H3K9me3 was abolished at all heterochromatic loci in both *cul4-K680R* and *cul4-1* cells (Fig 6B). Interestingly, H3K9me3 was lost even at loci where *cul4* mutations allowed FLAG-Clr4 localization – such as the pericentromeric *dg* and *dh* repeats with increased FLAG-Clr4 association in *cul4-1* cells, and the rDNA repeats where FLAG-Clr4 localization was unaffected by *cul4* mutations. These results suggest that Clr4 recruitment alone is insufficient for H3K9 methylation in the absence of Cul4 activity. These findings were further validated by ChIP-qPCR (Fig 6C-N and S6A-F). ChIP-qPCR showed that FLAG-Clr4 showed increased binding to centromeric *dg* and *dh* repeats in *cul4-1* cells relative to wild-type, whereas its binding to the mating type locus and telomeric region was reduced (Fig 6C-E and S6A). In *cul4-K680R* cells, FLAG-Clr4 binding was dramatically reduced at all tested loci. Consistent with the ChIP-seq data, H3K9me3 signal was lost at all heterochromatic loci in both *cul4* mutants (Fig 6E-G and S6B). Together, these results indicate that Cul4-mediated ubiquitination is required downstream of Clr4 recruitment to activate H3K9 methylation and facilitate Clr4 spreading beyond nucleation sites.

We next examined the effects of H3K14 substitutions on H3K9 methylation and FLAG-Clr4 recruitment to heterochromatic loci. Both H3K14R and H3K14A substitutions, which block ubiquitination, showed a complete loss of H3K9me2 and H3K9me3 at the pericentromeric *dg* and *dh* repeats, the mating type (*mat2P*) locus, and the subtelomeric *tlh1* gene, reducing levels to those observed in *clr4Δ* cells (Fig 6F-N, Fig S6C-H). Similarly, the localization of FLAG-Clr4 to each of these regions was also reduced to near background levels in both H3K14 mutant backgrounds (Fig 6J, M; Figure S6E, H). Together, these results suggest that H3K14 ubiquitination is required for Clr4 recruitment to heterochromatic regions and H3K9 methylation.

To assess the possible contribution of Clr4 ubiquitination to heterochromatin formation, we examined the effect of lysine-to-arginine mutations in Clr4 that impair its ubiquitination (Clr4 1^st^ half KtoR, Fig 5D). Clr4 1^st^ half KtoR mutations had little or no effect on silencing of *ade6*^*+*^ reporter genes located near strong H3K9me nucleation sites – *ort1R::ade6*^*+*^ and *10xtetO::ade6*^*+*^, but impaired the silencing at the *mat2P::ade6*^*+*^ reporter, positioned away from nucleation centers (Fig S7A-C). These findings suggest that Clr4 ubiquitination is dispensable for silencing near robust nucleation sites, such as those governed by RNAi-dependent or TetR-Clr4-mediated nucleation, but is important for silencing at loci where H3K9 methylation depends on Clr4 spreading. Consistent with these observations, Clr4 1^st^ half KtoR mutations did not affect H3K9me2, H3K9me3, or FLAG-Clr4 levels at the centromeric *dg* and *dh* repeats, but significantly reduced FLAG-Clr4 levels at the mating type (*mat2P*) and telomeric *tlh1* loci (Fig 6K, N and S6C-H). Despite the reduction in Clr4 association, H3K9me3 levels at *mat2P* were only modestly decreased, and H3K9me2 levels remained unchanged (Fig 6L, M). These results suggest that Clr4 ubiquitination enhances spreading and silencing at loci distal to nucleation sites, likely by promoting efficient read-write propagation of H3K9 methylation. In contrast, loss of H3K14 ubiquitination abolishes silencing entirely, underscoring its essential role in Clr4 recruitment and activation.

## Discussion

In this study, we show how the ubiquitination and methyl-transferase activities of the CLRC complex regulate histone H3K9 methylation and heterochromatin formation. Our findings suggest that Clr4-mediated H3K9 methylation is regulated at multiple levels. First, Clr4 itself serves a critical role in H3 ubiquitination with its active site serving as a substrate receptor for the specific ubiquitination of H3K14 by the Cul4 E3 ubiquitin ligase subunit of CLRC (Fig 7A, left). Second, while H3K14 ubiquitination strongly activates Clr4 to methylate H3K9 on the same H3 tail in cis (Fig 7B), Clr4 automethylation is required for Clr4 to methylate K9 on an unmodified H3 in trans (Fig 7C). Third, Cul4 ubiquitinates Clr4 leading to its release from the CLRC complex in a step that is required for the spreading of H3K9 methylation (Fig 7A, right). These results combined with structural modeling suggest that H3K14 ubiquitination controls Clr4 activity via 2 distinct interaction sites on Clr4. Ubiquitin interaction site 1 (UBS1) involves interactions between ubiquitin and the catalytic SET domain and positions H3K9 in the active site for cis methylation (Fig 7B). Ubiquitin interaction site 2 (UBS2), identified here, involves H3K14ub interactions with the chromodomain and ARL of Clr4, and coupled with Clr4 automethylation, licenses Clr4 for trans methylation of K9 on an unmodified H3 tail (Fig 7C). H3K14 ubiquitination therefore regulates both the substrate recognition (read) and catalytic activity (write) of Clr4. Below we discuss the implications of these findings for mechanisms that promote the highly selective ubiquitination of the flexible H3 amino terminus and how enzymatic activity is regulated to limit inappropriate heterochromatin formation.

**Figure 7.**
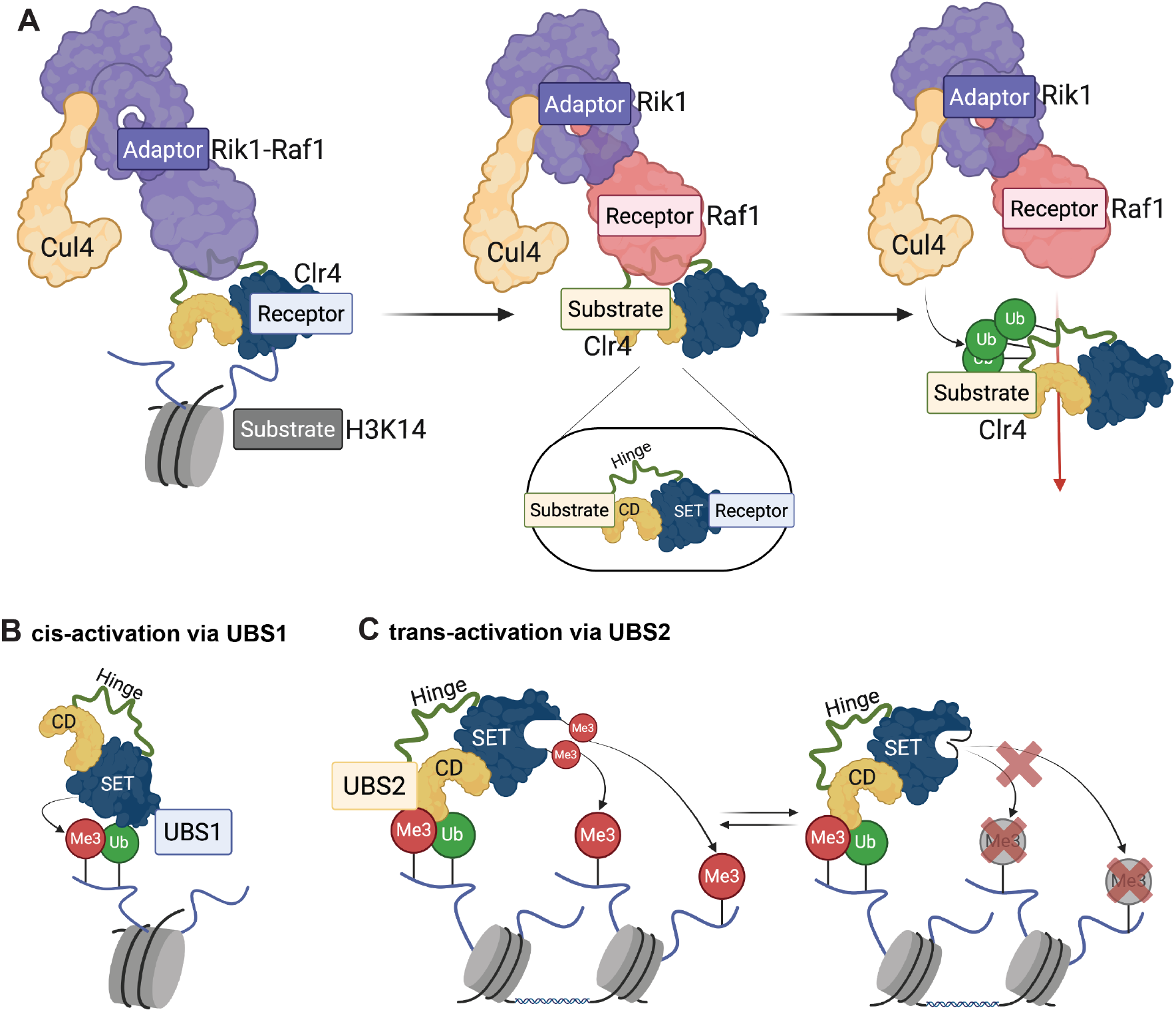
H3K14 ubiquitination regulates both the substrate recognition (“read”) and catalytic activity (“write”) of Clr4. **(A)** Clr4 as a dual substrate receptor within the CLRC complex. Left: The catalytic SET domain of Clr4 acts as a substrate receptor for histone H3, enabling Cul4mediated monoubiquitination of H3K14. Middle: The N-terminus of Clr4 interacts directly with Raf1 that functions as a substrate receptor for Clr4 ubiquitination on multiple surface-exposed lysines, predominantly located within the hinge region of Clr4. Right: Cul4-mediated ubiquitination of Clr4 promotes its dissociation from the CLRC complex, a step required for the spreading of H3K9 methylation. **(B)** Ubiquitin-binding site 1 (UBS1) drives cis methylation. H3K14 monoubiquitination strongly stimulates Clr4 to methylate H3K9 on the same histone tail in cis. This is mediated by UBS1, an interaction surface between ubiquitin and the SET domain that positions H3K9 in the catalytic site. **(C)** Ubiquitin-binding site 2 (UBS2) and automethylation enable trans methylation. UBS2, identified in this study, involves interactions between H3K14ub and the chromodomain-ARL interface within Clr4. Together with Clr4 automethylation of the ARL, UBS2 licenses Clr4 to methylate H3K9 on an unmodified histone tail in trans, thereby facilitating methylation spreading.

### Clr4 acts as the substrate receptor for Cullin 4-mediated ubiquitination

The highly flexible amino terminus of H3 contains lysines at positions 4, 9, 14, 18, and 23. The highly selective ubiquitination of H3K14 by the CLRC complex suggests that the substrate receptor subunit of the complex must present the H3 amino terminus so that K14 is preferentially ubiquitinated. The presumed substrate receptor of the CLRC complex, Raf1, which is a homolog of DDB2 in other CRL4 complexes, lacks an obvious histone tail binding activity. Instead, our findings demonstrate that the catalytic pocket of the Clr4 methyltransferase serves a substrate receptor for Cul4-mediated ubiquitination of H3K14. The unmodified H3 amino terminus may bind to a partially open Clr4 active site so that K14 is presented to Cul4-E2 for ubiquitination. Following K14 ubiquitination, K9 becomes optimally located in the catalytic pocket and is efficiently methylated. Our results show that relative to nucleosomes containing unmodified H3K9, H3K9me3 nucleosomes are poor substrates for Cul4-mediated ubiquitination, suggesting that K14 ubiquitination occurs prior to K9 methylation and the release of the H3 tail from the Clr4 active site. In principle, the chromodomain of Clr4 may also be able to present the H3 tail for K14 ubiquitination, but we found that the chromodomain was dispensable for H3K14 ubiquitination. The apparent conservation of ubiquitin binding sites in the mammalian H3K9 methyl-transferases, SETDB1, SUV39H1, and SUV39H2 (Fig S8), and the reported coupling of H3K9 methylation and H3K14 and K18 ubiquitination ^13,14^ suggest a conserved and fundamental role for the coupling of H3 ubiquitination and methylation and raise the possibility that the active sites of H3K9 methyltransferases play conserved roles as substrate receptors for E3 ubiquitin ligases that activate them.

### A key role for H3K14ub in both the read and write functions of Clr4

A key outcome of our reconstitution experiments is the demonstration that an initial ubiquitination event on one H3 amino terminal tail allows the trans methylation of an unmodified H3 tail. This finding suggests that the association of Clr4 with H3K14ub promotes an open active site conformation that promotes trans methylation. Additionally, they predict the existence of a second ubiquitin binding site that mediates this activation. Consistent with this expectation, the chromodomain exerts an inhibitory effect on Clr4 methyltransferase activity and our XL-MS and structural modeling experiments suggest that the chromodomain and ARL form a second ubiquitin binding site, UBS2, which together with Clr4 automethylation allows trans-methylation. Our observation that the chromodomain is required for the association of Clr4 with ubiquitinated nucleosomes further supports this model. Ubiquitination is therefore likely to control both the read and write functions of Clr4 via a “ubiquitination activation cycle”. In this cycle, the chromodomain reads both H3K9me3 and H3K14ub leading to the relief of its inhibitory effect on the SET domain. This then leads to Clr4 automethylation and the trans methylation and ubiquitination of an unmodified H3 tail. Repeated cycles of H3 methylation and ubiquitination then lead to the spreading of heterochromatin beyond nucleation sites. In support of this model, substitution of H3K14 with arginine, abolishes H3K9 methylation and Cul4 mutations that block its ubiquitination activity similarly block H3K9 methylation. However, in these ubiquitination-defective cells, we observed weak association of Clr4 with RNAi nucleation sites, indicating that ubiquitination is not required for the initial recruitment of the CLRC complex.

### The relative contributions of H3K14 and Clr4 ubiquitination

Our findings suggest that the Clr4 and H3K14 ubiquitination events play distinct roles in heterochromatin formation. H4K14 ubiquitination is critical for activation of Clr4 methyltransferase activity and H3K9 methylation, largely independently of Clr4 recruitment to nucleation sites. Thus, Clr4 recruitment in the absence of H3K14 ubiquitination, as in *cul4* mutant cells, does not lead to detectable H3K9 methylation. Disruption of Clr4 ubiquitination, on the other hand, has little or no effect on H3K9 methylation or reporter gene silencing at strong RNAi nucleation sites, such as the centromeric dg/dh and the telomeric tlh1 loci, but results in diminished Clr4 recruitment and H3K9me3 at the mating type locus where heterochromatin formation can occur independently of RNAi. Therefore, while H3K14 ubiquitination is an obligatory step in H3K9 methylation and heterochromatin formation, Clr4 ubiquitination appears to play a regulatory role that may promote the spreading of H3K9me3 beyond nucleation sites. Mechanistically, our observation that Clr4 ubiquitination promotes its release from the CLRC complex suggests that redistribution of Clr4 from strong nucleation sites to other regions may regulate heterochromatin dynamics, and may be related to the more efficient incorporation of ubiquitinated Clr4 into Swi6 condensates ^11^.

## Acknowledgements

We thank the Harvard Cryo-Electron Microscopy Center for Structural Biology for training and access to cryo-EM microscopes, members of the Moazed and Farnung labs, Reza Behrouzi, Vaclav Veverka, Rudi Pisa, Mark Currie, and Nahid Iglesias for helpful discussions, and Vaclav Veverka for comments on the manuscript and assistance with YASARA and COOT structural modeling. This work was supported by an EMBO long-term fellowship LTF-451 (K.P.), NIH R44 GM119893 (EpiCypher), and NIH RO1 GM072805 (D.M.). L.F. is an HHMI Freeman Hrabowski Scholar and D.M. is an HHMI Investigator.

## Author contributions

K.P. and D.M. designed experiments. K.P. performed all experiments except as follows: S.S.P. performed strain construction for ChIP-seq, ChIP-qPCR, and silencing assays, and carried out the experiments shown in Fig 6, S6, and S7. J.A.P., X.L., and M.K. performed mass spectrometry. M.K., S.S.P., and K.P. constructed strains for Fig 5 and S5. M.K. performed the initial characterization of Clr4–Ub. E.F.P., R.W., and M.A.C. synthesized the H3K14ub nucleosomes, and M.C.K. (EpiCypher) served as project manager for H3K14ub nucleosome preparation. L.F., S.P.G., and D.M. supervised research. K.P. and D.M. wrote the manuscript with input from all authors.

## Materials and Methods

### Strains and plasmids construction

*S. pombe* strains used in this study and their genotypes are listed in Table S2. Plasmids are listed in Table S3.

The coding sequence of *S. pombe* Rik1, Raf1, Raf2 and Cul4 were amplified from cDNA. Using ligation-independent cloning, Rik1, Raf2 and Cul4 was cloned into 438-A vector (Addgene: 55218). Raf1 was cloned into 438-C vector (Addgene: 55220) containing an N-terminal 6×His tag followed by a maltose binding protein (MBP) tag and a tobacco etch virus protease (TEV) cleavage site. The plasmids were subsequently combined using ligation-independent cloning to integrate multiple genes flanked by PolH promoter and SV40 terminator into the AcMNPV genome.

The plasmids pGEX-6P-1-Clr4 (GST-Clr4 WT, pDM2113), pGEX-6P-1-Clr4 192-490 (GST-Clr4 ΔCDΔhinge, alternatively GST-Clr4 SET, pDM1906), pGEX-6P-1-Clr4-Y451N (GST-Clr4^dead^, pDM2142), pGEX-6P-1-Clr4-K455,472R (GST-Clr4 K455,472R, pDM2132) used in this study were previously described in ^8^. pGEX-6P-1-Clr4-W31G (GST-Clr4 W31G, pDM2433) and pGEX-6P-1-Clr4-W31G,Y451N (GST-Clr4 W31G,Y451N) was generated by introducing the tryptophan-to-glycine mutation to a vector encoding wild-type Clr4 and Clr4^dead^, respectively, using QuikChange II Site-Directed Mutagenesis Kit from Agilent Technologies. The Clr4 truncations were generated by Quik-Change II Site-Directed Mutagenesis Kit using GST-Clr4 WT, with the Clr4 sequence encoding amino acids 1-69 (GST-Clr4 ΔhingeΔSET, pDM2432), the Clr4 sequence encoding amino acids 70-490 (GST-Clr4 ΔCD, pDM1000), and the Clr4 sequence encoding amino acids 1-192 (GST-Clr4 ΔSET, pDM2426). Clr4 Δhinge was generated by introducing either a short six amino acid flexible linker (GSSGSS, GST-Clr4 Δhinge-s) or a long 18 amino acid flexible linker (6 repeats of GSS sequence, GST-Clr4 Δhinge-l) in place of the hinge region, amino acids 70-191, using site-directed mutagenesis. For the plasmid pGEX-6P-1 GST-Clr4 CD-Hinge-Set2 (GST-Clr4N-Set2, pDM2428), the amino acid sequence 90-330 of *S. pombe* SET2 was amplified from genomic DNA and fused to the C-terminus of GST-Clr4 ΔSET. All plasmid sequences were verified by DNA sequencing.

The H3K9M, H3K14R and H3K9M,K14R variants were generated by introducing the single amino acid mutation to a vector containing H3C110A using QuikChange II Site-Directed Mutagenesis Kit.

*S. pombe* strains were generated by either transformation of linear DNA using the lithium acetate transformation protocol ^24^ or by genetic crosses and random spore analysis. Integrations were screened by antibiotic markers and confirmed by colony PCR.

### Protein expression and purification

The expression of CLRC^-Clr4^ complex in Hi5 insect cells was performed as previously described in ^25^. All insect cell lines were maintained at 1×10^6^ cells/mL, and cell viability was monitored to remain above 85% by periodic supplementation with fresh medium. Cells were harvested by centrifugation (300×*g*, 4 °C, 25 min) and resuspended in lysis buffer [50 mM HEPES pH 7.5, 500 mM NaCl, 1 mM TCEP, 1 mM PMSF, 30 mM imidazole, 10% glycerol, protease inhibitors]. Cell pellets were lysed using a manual Dounce tissue grinder (B-insert, 2 × 20 strokes in 40 mL total volume, Sigma-Aldrich). The lysate was clarified by centrifugation (20 min, 23,000×*g*, 4 °C) and loaded onto Ni-NTA resin (NEB) and washed with buffer E [50 mM HEPES pH 7.5, 300 mM NaCl, 50 mM imidazole, 1 mM MgCl_2_, 10% glycerol, 1 mM TCEP, 1 mM PMSF, protease inhibitors]. Elution was performed using buffer F [buffer E with 0.5 M imidazole]. The eluate was then incubated with amylose resin (NEB) for 20 min, washed with buffer M [50 mM HEPES pH 7.5, 300 mM NaCl, 1 mM MgCl_2_, 10% glycerol, 1 mM TCEP, 1 mM PMSF, protease inhibitors], and eluted with the same buffer supplemented with 50 mM maltose. Eluate was concentrated using a 50 kDa MWCO Amicon centrifugal filter unit to ~0.5 mL. Size-exclusion chromatography (SEC) was performed using a Superose 6 Increase 10/300 column equilibrated in SEC buffer [20 mM HEPES pH 7.5, 300 mM NaCl, 1 mM TCEP, 1 mM MgCl_2_, 10% glycerol].

The expression and purification of all Clr4 mutants was performed as described previously ^8^. The final SEC was performed using a Superdex 200 Increase 10/300 column equilibrated in SEC buffer [20 mM HEPES pH 7.5, 300 mM NaCl, 1 mM TCEP, 1 mM MgCl_2_, 10% glycerol]. All *Xenopus laevis* histones were expressed and purified as described previously ^26,27^. Nucleosome reconstitution using the H3K9M, H3K14R and H3K9M,K14R variants was performed using a salt-gradient dialysis as described previously ^26,27^. Nucleosomes were subsequently purified by SEC using a Superose 6 Increase 10/300 column equilibrated in SEC buffer [20 mM HEPES pH 7.5, 20 mM NaCl, 1 mM TCEP, 10% glycerol] and concentrated using a 50 kDa MWCO Amicon centrifugal filter unit.

Methylated (H3K9me3) and/or ubiquitinated (H3K14ub) nucleosomes are available from EpiCypher (SKU: 16-0315 and 16-0398, respectively). A double-modified nucleosome harboring H3K9me3K14ub was custom-prepared by EpiCypher for this study. All EpiCypher nucleosomes were assembled through salt-dialysis of semi-synthetic histones with 5’ biotinylated DNA (147bp of 601 nucleosome positioning sequence) as described previously ^28,29^ and validated by SDS-PAGE, immunoblotting, and mass spectrometry (as appropriate). All ubiquitinated histones contain a native gamma-lysine isopeptide linkage cleavable by deubiquitinating (DUB) enzymes^30^.

### GST pull-down assays

Pulldown assays were performed using 10 µL of glutathione Sepharose beads (Cytiva) for GST-tagged proteins or 100 µL of amylose resin (NEB) for MBP-tagged proteins. Reaction components were mixed at a 1:1 molar ratio and incubated on ice for 30 minutes. For Clr4 ubiquitination assays, reactions were carried out at 30 °C for 2 hours prior to bead incubation. Following incubation, 5 µL of each reaction was reserved as input, and the remaining volume was incubated with the appropriate affinity resin at 4 °C for 1 hour. Beads were washed three times with wash buffer [20 mM HEPES pH 7.5, 150 mM NaCl, 1 mM TCEP, 1 mM MgCl_2_, 10% glycerol, 0.05 mg/mL insulin, 0.02% Triton X-100], and bound proteins were eluted using 0.5 M ammonium hydroxide (NH_4_OH). Eluates were dried by SpeedVac and analyzed by SDS-PAGE.

### Isolation of ubiquitinated proteins

*S. pombe* cells were transformed with pREP1-nmt1-LEU2 encoding 6×His-tagged ubiquitin (6×His-Ub) ^31^ using the lithium acetate method ^24^. Transformants were selected on EMM lacking leucine, and overexpression was induced by omitting thiamine from the medium. Cells (200 OD) were harvested (10 min, 3500×*g*, 4 °C), washed in ddH_2_O (5 min, 2500×*g*, 4 °C), and flash frozen in liquid nitrogen. Lysis was performed with 6 mL 1.91 N NaOH and 7.5% β-mercaptoethanol on ice (15 min), followed by precipitation with an equal volume of 55% TCA (15 min, on ice). Lysed cells were pelleted by centrifugation (15 min, 2500×*g*, 4 °C), washed twice with ice-cold acetone, and resuspended in 12 mL buffer A [6 M guanidinium chloride, 100 mM NaH_2_PO_4_, 10 mM Tris-HCl pH 8.0, 0.05% Tween-20]. After 1 h shaking at RT, lysates were clarified by centrifugation (20 min, 23,000×*g*, 4 °C), supplemented with 20 mM imidazole, and incubated overnight with 100 µL Ni-NTA Magnetic Agarose beads (Qiagen) at 4 °C. Beads were washed 3× with buffer A + 20 mM imidazole and 5× with buffer C [8 M urea, 100 mM NaH_2_PO_4_, 10 mM Tris-HCl pH 6.3, 0.05% Tween-20], and eluted with 30 µL 1% SDS (10 min, 65 °C). Eluates were dried in a SpeedVac (45 °C, ~25 min), resuspended in 10 µL ddH_2_O and 15 µL HU buffer [8 M urea, 5% SDS, 200 mM Tris-HCl pH 6.8, bromophenol blue, 1.5% DTT], and incubated for 10 min at 65 °C before analysis on SDS-PAGE gel.

### Co-Immunoprecipitation (Co-IP)

*S. pombe* cells (OD_600_ = 200) in logarithmic phase were harvested by centrifugation (6000 rpm, 10 min, 4 °C), washed twice with cold 1× TBS, and resuspended in one-fifth volume of lysis buffer [20 mM HEPES pH 7.5, 100 mM NaCl, 5 mM MgCl_2_, 1 mM EDTA, 10% glycerol]. Cells were frozen in liquid nitrogen and disrupted using a Freezer Mill (12 cycles, 1 min 45 s grinding, 2 min cooling, 10 CPS). Lysates were treated with 4 µL Benzonase (250 U/µL) per mL lysate at 4 °C for 1 h, then clarified by centrifugation (13,200 rpm, 15 min, 4 °C). Supernatants were used directly for binding. Protein A or Protein G magnetic beads (20 µL per sample) were pre-washed in lysis buffer and incubated overnight at 4 °C with antibody in lysis buffer. Crosslinking was performed using 3.84 mg/mL DMP in 0.2 M sodium borate (pH 9.0) for 30 min at room temperature, followed by quenching with 0.2 M ethanolamine (pH 8.0, 1.5 h). Beads were washed with wash buffer (lysis buffer containing 0.25% Triton X-100, 0.5 mM DTT, 1 mM PMSF and protease inhibitor cocktail) and added to clarified lysates. Immunoprecipitations were carried out for 3 h at 4 °C with rotation. Beads were washed three times with wash buffer. For mass spectrometry samples, detergent was omitted and washes included one round in wash buffer followed by two rounds in 1× TBS. Proteins were eluted with 50 µL of 500 mM ammonium hydroxide (20 min, 37 °C). Eluates were split for western blot (30 µL) and mass spectrometry (20 µL).

### Methyltransferase assays

The methyltransferase assay was performed as previously described ^8^, with minor modifications. Reactions were carried out using 1 µM reconstituted nucleosomes assembled with *Xenopus laevis* histones containing wild-type H3 or mutant variants (H3K9M, H3K14R, or H3K9M,K14R). Where indicated, methylated (H3K9me3) and/or ubiquitinated (H3K14ub) nucleosomes used as substrates were purchased from EpiCypher (SKU: 16-0315 and 16-0398, respectively) ^28– 30,32,33^. Reaction products were resolved by SDS-PAGE, and the gel was scanned and incubated for 90 minutes in EN3HANCE™ autoradiography enhancer (Perkin Elmer). Following incubation, the gel was rinsed with water and dried using a gel dryer for 90 minutes. Incorporation of [^3^H]-methyl groups was visualized by exposing the dried gel to Amersham Hyperfilm at −80 °C for 2 to 72 hours.

### Ubiquitination assays

Ubiquitination reactions were carried out in a total volume of 20 µL containing 1 µM reconstituted nucleosomes or 10 µM histone H3 N-terminal peptide (aa 1–20; AnaSpec) as substrates, 2 µM Clr4, and 1 µM CLRC^-Clr4^ complex. The reconstituted nucleosomes were assembled with *Xenopus laevis* histones containing wild-type H3 or mutant variants (H3K9M, H3K14R, or H3K9M,K14R) and methylated (H3K9me3) nucleosomes were purchased from EpiCypher (SKU: 16-0315) ^28–30,32,33^. Reaction components excluding the ubiquitination mix were pre-incubated on ice for 30 minutes. A 5× ubiquitination mix was prepared in reaction buffer [50 mM Tris-HCl pH 7.5, 2.5 mM MgCl_2_, 0.5 mM DTT] and contained 500 nM recombinant human E1 (UBE1), 1 µM recombinant E2 (UbcH5c/UBE2D3), 20 mM ATP, and 50 µM recombinant human HA-Ubiquitin (all from R&D Systems, Bio-Techne) together with 100 µM S-adenosylhomocysteine (SAH). The ubiquitination mix was pre-incubated at 37 °C for 20 minutes prior to use. Final reactions containing 1× ubiquitination mix were incubated at 30 °C for 15 minutes with continuous shaking at 300 rpm. Reactions were quenched by adding 5 µL of 5× SDS loading buffer (Fisher Scientific) and prepared for SDS-PAGE analysis.

Combined methyltransferase–ubiquitination reactions were performed in reaction buffer [50 mM Tris-HCl pH 7.5, 2.5 mM MgCl_2_, 0.5 mM DTT] containing 1× ubiquitination mix as described above, with the exception that S-adenosylhomocysteine (SAH) was replaced by 0.42 µM [^3^H]-S-adenosyl methionine ([^3^H]-SAM; Perkin Elmer). Reactions were incubated at 30 °C for 30 minutes with continuous shaking at 300 rpm and processed according to the methyl-transferase assay protocol.

### Cross-linking Mass Spectrometry (XL-MS)

Cross-linking mass spectrometry (XL-MS) was performed as previously described ^34,35^, with the following modifications. To prepare the cross-linked sample, 100 µL of 2 µM H3K14ub nucleosome complex was incubated with a 7-fold molar excess of Clr4 (14 µM) in the presence of 135 µM S-adenosylhomocysteine (SAH) for 1 hour on ice. Recombinant nucleosomes containing monoubiquitinated histone H3K14 (H3K14ub) were purchased from EpiCypher (SKU: 16-0398). The complex was purified by size-exclusion chromatography using a Superose 6 Increase 3.2/300 column (Cytiva) on an ÄKTA Micro FPLC system. Fractions corresponding to the complex were pooled (~1 µM, 70 µL), and freshly prepared BS3 (Bis[sulfosuccinimidyl] suberate; 10 mM in HEPES buffer, pH 7.8) was added to a final concentration of 1 mM. Cross-linking was carried out at room temperature (RT) for 1 hour with gentle agitation. The reaction was quenched by addition of hydroxylamine (final concentration 100 mM) and incubated for 15 minutes at RT with mixing. Benzonase (250 units, Sigma Aldrich) was then added, and samples were incubated for 30 minutes at 4 °C. Proteins were denatured by the addition of 8 M urea, reduced with 5 mM TCEP (15 minutes, RT), alkylated with 10 mM iodoacetamide (IAA; 30 minutes, in the dark), and quenched with 5 mM DTT (15 minutes, RT). Proteins were precipitated by adding trichloroacetic acid (TCA) to 20%, followed by incubation on ice for 30 minutes. Precipitates were collected by centrifugation at 14,000 rpm for 15 minutes at 4 °C, washed once with 1 mL ice-cold acetone, and once with 1 mL methanol. After each wash, samples were centrifuged at 14,000 rpm for 5 minutes at 4 °C. Pellets were resuspended in 200 mM EPPS, pH 8.5, and digested at room temperature for 14 h with LysC protease at a 100:1 protein-to-protease ratio. Trypsin was then added at a 100:1 protein-to-protease ratio and the reaction was incubated for 6 h at 37 °C. Digested peptides were acidified with 10% formic acid to pH ~2, desalted using stage tips with Empore C18 SPE Extraction Disks (3M), and dried under vacuum. The sample was reconstituted in 5% formic acid (FA)/5% acetonitrile and analyzed in the Orbitrap Lumos Mass Spectrometer (Thermo Fisher Scientific) coupled to an EASY-nLC 1200 (Thermo Fisher Scientific) ultra-high-pressure liquid chromatography (UHPLC) pump, as well as a high-Field Asymmetric waveform Ion Mobility Spectrometry (FAIMS) interface. Peptides were separated on an in-house packed 100 µm inner diameter column with 35 cm of Accucore C18 resin (2.6 µm, 150 Å, Thermo Fisher), using a gradient of 5–35% acetonitrile (0.125% FA) over 135 min at ~500 nL/min. The instrument was operated in data-dependent mode. FTMS1 spectra were collected at a resolution of 120K, with an automatic gain control (AGC) target of 5 × 10^5^ and a maximum injection time of 50 ms. The most intense ions were selected for MS/MS for 1 s in top-speed mode, while switching among three FAIMS compensation voltages (CV): −40, −60, and −80 V in the same method. Precursors were filtered by charge state (allowed: 3 ≤ z ≤ 7), and monoisotopic peak assignment was turned on. Previously interrogated precursors were excluded using a dynamic exclusion window (60 s ± 7 ppm). MS2 precursors were isolated with a quadrupole mass filter set to a width of 0.7 m/z and analyzed by FTMS2, with the Orbitrap operating at 30K resolution, an AGC target of 100K, and a maximum injection time of 150 ms. Precursors were fragmented by high-energy collision dissociation (HCD) at 30% normalized collision energy.

Mass spectra were processed and searched using the PIXL search engine ^34^. The sequence database contained proteins identified at 1% FDR in a non-cross-linked Comet search ^36^. For the PIXL search, precursor tolerance was set to 15 ppm and fragment ion tolerance to 10 ppm. Searches included methionine oxidation and protein N-terminal acetylation as variable modifications, as well as relevant monolink masses-Lys modification of 156.0786 for BS3. Crosslinked peptides were searched assuming BS3 crosslinker +138.0681. Searches considered 60 protein sequences to ensure sufficient statistics for FDR estimation. Matches were filtered to 1% FDR at the unique peptide level using linear discriminant features as previously described ^34^.

### K-GG Mass Spectrometry for Ubiquitination Site Identification

Ubiquitination reactions were performed in a final volume of 20 µL containing 2 µM Clr4 and 1 µM CLRC-Clr4 complex. All components except the ubiquitination mix were pre-incubated on ice for 30 minutes. A 5× ubiquitination mix was prepared in reaction buffer [50 mM Tris-HCl (pH 7.5), 2.5 mM MgCl_2_, 0.5 mM DTT], and included 500 nM recombinant human E1 (UBE1), 1 µM E2 enzyme (UbcH5c/UBE2D3), 20 mM ATP, 50 µM recombinant HA-tagged ubiquitin, and 100 µM S-adenosylhomocysteine (SAH) (all reagents from R&D Systems, Bio-Techne). The ubiquitination mix was pre-activated at 37 °C for 20 minutes before use. Final reactions containing 1× ubiquitination mix were assembled and incubated at 30 °C for 30 minutes with continuous shaking at 300 rpm. Proteins were precipitated by adding trichloroacetic acid (TCA) to 20%, followed by incubation on ice for 30 minutes. Precipitates were collected by centrifugation at 14,000 rpm for 15 minutes at 4 °C, washed once with 1 mL of ice-cold acetone, and once with 1 mL of methanol. After each wash, samples were centrifuged at 14,000 rpm for 5 minutes at 4 °C.

Mass spectrometry data were collected using an Orbitrap Astral mass spectrometer (Thermo Fisher Scientific) coupled with a Neo Vanquish liquid chromatograph. Peptides were separated on a 110 cm µPAC C18 column (Thermo Fisher Scientific). For each analysis, ~0.5 µg was loaded onto the column. Peptides were separated using a 75 min gradient of 5–29% acetonitrile in 0.125% formic acid with a flow rate of 300 µL/min. The scan sequence began with an Orbitrap MS1 spectrum with the following parameters: resolution 60,000, scan range 350–1350 Th, AGC target 200%, maximum injection time 50 ms, RF lens setting 50%, and centroid spectrum data type. FAIMS was enabled using the Top20 setting for each run. Samples were analyzed three times with two CV sets: −35 V, −45 V, −55 V, −60 V, and −70 V (for two replicates), and −40 V, −60 V, −80 V (for one replicate). Astral data acquisition included AGC 100%, maximum injection time 25 ms, isolation window 1.2 Th, normalized collision energy (NCE) 28, and centroid spectrum data type. Unassigned and singly charged species were excluded from MS2 analysis, and dynamic exclusion was set to 15 s across CVs.

Mass spectrometric data analysis was performed using a Comet-based in-house software pipeline. MS spectra were converted to mzXML using MSConvert. Database searching included the protein of interest plus common contaminants, concatenated with a reverse database of all sequences in reverse order. The digest was set to non-specific. Searches were performed with a 3 Da precursor ion tolerance and a product ion tolerance of 0.02 Th. Oxidation of methionine residues (+15.9949 Da) and ubiquitylation (+114.0429 Da) at lysine residues were set as variable modifications. PSM filtering was performed manually, considering only tryptic peptides, an XCorr > 1.5, and an absolute PPM mass tolerance <10. Spectra were manually validated.

### Mass Spectrometry Analysis of Purified Complex Components

Purified protein complexes were precipitated by adding trichloroacetic acid (TCA) to 20%, followed by incubation on ice for 30 minutes. Precipitates were collected by centrifugation at 14,000 rpm for 15 minutes at 4 °C, washed once with 1 mL of ice-cold acetone, and once with 1 mL of methanol. Pellets were resuspended in 200 mM EPPS, pH 8.5, and digested at room temperature for 14 h with LysC protease at a 100:1 protein-to-protease ratio. Trypsin was then added at the same ratio, and the reaction was incubated for 6 h at 37 °C.

Mass spectrometric data were collected on an Orbitrap Lumos mass spectrometer coupled to an easyNano LC. Approximately 1 µg of peptide was separated at a flow rate of 350 nL/min on a 100 µm capillary column packed with 35 cm of Accucore 150 resin (2.6 µm, 150 Å; Thermo Fisher Scientific). The scan sequence began with an MS1 spectrum (Orbitrap analysis, resolution 60,000, 350–1350 Th, AGC target 100%, maximum injection time set to “auto”). Data were acquired over 90 minutes per fraction. The hrMS2 stage involved fragmentation by HCD (30% NCE) and Orbitrap analysis (AGC 200%, injection time 60 ms, isolation window 1.2 Th, resolution 7,500). Data were acquired using the FAIMSpro interface with a dispersion voltage (DV) of 5,000 V. The sample was analyzed three times with three CV sets: set 1 (−30 V, −50 V, −70 V), set 2 (−40 V, −60 V, −80 V), and set 3 (−45 V, −55 V, −65 V). The TopSpeed parameter was set to 1 s per CV. Unassigned and singly charged species were excluded from MS2 analysis, and dynamic exclusion was set to 90 s across CVs.

Mass spectra were processed using a Comet-based inhouse software pipeline. MS spectra were converted to mzXML using MSConvert. Database searches included all entries from the *S. pombe* UniProt database concatenated with a reverse database. Searches used a 50 ppm precursor ion tolerance and a 0.03 Th product ion tolerance. Carbamidomethylation of cysteine residues (+57.0215 Da) was set as a static modification, while oxidation of methionine (+15.9949 Da) was set as a variable modification.

Peptide spectral matches (PSMs) were filtered to 1% FDR ^37,38^. PSM filtering was performed using linear discriminant analysis ^39^, considering parameters including XCorr, ΔCn, missed cleavages, peptide length, charge state, and precursor mass accuracy. PSMs were identified, quantified, and collapsed to 1% FDR, then further to a final protein-level FDR of 1%. Protein assembly followed the principle of parsimony. Spectral counts were extracted and analyzed.

### Immunoprecipitation and Mass Spectrometry Analysis

For mass spectrometry, bound proteins were eluted from the affinity matrix using 0.5 M ammonium hydroxide and subsequently dried to completion in a vacuum centrifuge. Dried eluates were reconstituted in 200 mM EPPS buffer (pH 8.5) and digested with sequencing-grade trypsin (Promega; cat. #V5111) in the presence of 2% (v/v) acetonitrile. Digestion was carried out overnight at 37 °C. Resulting peptides were directly labeled with TMT16plex reagents (Thermo Fisher Scientific; cat. #A44520) following the manufacturer’s instructions. Labeling efficiency was verified by LC-MS/MS. Reactions were quenched with 0.3% (v/v) hydroxylamine for 10 minutes, pooled, acidified with formic acid, and dried to near completion using a speed vacuum concentrator.

Peptides were fractionated by high-pH reversed-phase chromatography using the Pierce High pH Reversed-Phase Peptide Fractionation Kit (Thermo Fisher Scientific; cat. #84868). Elution was performed stepwise using increasing concentrations of acetonitrile: 10%, 12.5%, 15%, 17.5%, 20%, 25%, 30%, 35%, 40%, 50%, 65%, and 80%. Fractions were concatenated into six pools (1+7, 2+8, 3+9, 4+10, 5+11, 6+12), dried under vacuum, desalted using StageTips, and subjected to LC-MS/MS.

Peptides were analyzed on an Orbitrap Lumos Tribrid mass spectrometer (Thermo Fisher Scientific) coupled to an EASY-nLC 1200 system (Thermo Fisher Scientific). Separation was performed using 100 µm inner diameter fused silica columns packed with 2.6 µm Accucore C18 resin (Thermo Fisher Scientific), using a linear gradient of acetonitrile in 0.1% formic acid.

MS1 spectra were acquired in the Orbitrap at 120,000 resolution across a mass range of 400–1400 m/z. Precursors were fragmented by collision-induced dissociation (CID) at 35% normalized collision energy, and MS2 spectra were collected in the ion trap. Synchronous precursor selection (SPS) was applied to isolate multiple MS2 fragment ions for further fragmentation by high-energy collision-induced dissociation (HCD) at 55% collision energy. TMT reporter ions were quantified in the Orbitrap at a resolution of 50,000 (at 200 m/z) using an MS3 scan.

Spectra were searched using an in-house pipeline based on SEQUEST (v.28, rev. 12), querying a combined forward and reversed human UniProt database (release: July 2014). Search parameters included a precursor mass tolerance of 50 ppm and a fragment ion tolerance of 0.9 Da. Up to two missed tryptic cleavages were allowed, and oxidation of methionine (+15.9949 Da) was included as a variable modification. Peptide-spectrum matches (PSMs) were filtered to a 1% false discovery rate (FDR) using a target-decoy approach combined with linear discriminant analysis (LDA). Protein-level FDR was also controlled at 1%. Quantification was based on the summed signal-to-noise (S/N) of TMT reporter ions, requiring a minimum summed S/N > 200 and an isolation specificity ≥ 70%.

Detailed procedures for TMT labeling, peptide cleanup, and fractionation have been described previously ^40^.

### Structure Prediction Using AlphaFold3

Protein structure predictions were generated using AlphaFold3 via the online interface at DeepMind and EMBLEBI’s AlphaFold Protein Structure Database (https://alphafold.ebi.ac.uk). Input sequences were submitted as FASTA files, and multimer predictions were enabled with default parameters. Predicted models were ranked using the internal scoring metrics based on the weighted combination of inter-chain predicted TM-score (ipTM) and pTM scores. The top-ranked models were selected for further analysis. Confidence in local structure and interaction elements was evaluated using per-residue pLDDT (predicted Local Distance Difference Test) and Predicted Aligned Error (PAE) plots. Structural visualization and inspection were performed using ChimeraX v1.9.

### Protein-Protein Docking Using HADDOCK

Protein-protein docking was performed using the High Ambiguity Driven protein-protein DOCKing (HADDOCK) web server (v2.4; https://wenmr.science.uu.nl/haddock2.4) ^41,42^. Input structures were prepared in PDB format and curated to remove heteroatoms and alternate conformations. Active and passive residues were defined based on crosslinking mass spectrometry (XL-MS) distance restraints and were limited to surface-exposed residues selected from models predicted by AlphaFold3. Docking proceeded in three stages: rigid-body energy minimization, semi-flexible refinement, and final refinement in explicit solvent. Default HADDOCK parameters were used unless otherwise specified. Resulting models were clustered based on RMSD with a cutoff of 7.5 Å, and ranked by HADDOCK score. The top-ranking cluster was used for downstream analysis. Structural visualization and interface analysis were performed using ChimeraX v1.9.

### Molecular Dynamics Simulations

Molecular dynamics (MD) simulations were performed using a protein structure provided in PDB format and processed with YASARA Structure (version ≥15.1.1) ^43^. To avoid introducing artificial terminal charges, the N-terminus was capped with an acetyl group, and the C-terminus was modified with an N-methyl amide group. The structure was confirmed to contain no disulfide bonds. All crystallographic waters, co-solvents, and ligands were removed prior to simulation. Protonation states were assigned at physiological pH 7.4 using YASARA’s built-in routines, which also optimized hydrogen bonding networks and completed missing side chains where applicable. The protein was placed in a cubic simulation box with a 10 Å buffer around the solute and solvated using explicit TIP3P water molecules. Sodium and chloride ions were added to neutralize the system and achieve a physiological ionic strength of 0.9% NaCl. A water density of 0.997 g/mL was used, corresponding to a simulation temperature of 298 K. Distance restraints were applied based on chemical crosslinking coupled with mass spectrometry (XL-MS). Crosslinked residue pairs, primarily involving lysines, were modeled as harmonic restraints between Cα atoms with equilibrium distances consistent with the span of the crosslinker (~25–30 Å). These restraints were incorporated prior to energy minimization and maintained throughout the simulation to preserve experimentally determined spatial proximities. Simulations were conducted using the AMBER14 all-atom force field, with an 8 Å cutoff for non-bonded interactions and Particle Mesh Ewald (PME) for long-range electrostatics. The system was simulated in the NPT ensemble, with temperature controlled via a rescaling thermostat and pressure maintained by solvent density scaling. Periodic boundary conditions were applied in all three spatial dimensions. A multiple time step integration scheme (2 × 1.25 fs = 2.5 fs total) was used. High-velocity atoms (>13,000 m/s) were automatically braked to ensure numerical stability. Before the production phase, energy minimization was carried out to relieve steric clashes and optimize geometry. The total simulation time was 10 ns, with snapshots saved every 100 ps. Simulations were performed using YASARA’s native trajectory format, which preserves both atomic coordinates and velocities. Built-in logic allowed for automatic resumption from previously saved snapshots in the event of an interruption. Trajectory analysis was carried out using YASARA’s internal tools. RMSD was monitored to assess overall structural stability, and the compliance of distance restraints was evaluated throughout the simulation.

### Cryo-EM

Cryo-EM sample was prepared by incubating 100 µL of 2 µM H3K14ub nucleosome complex with a 7-fold molar excess of Clr4 (14 µM) in the presence of 135 µM S-adenosylhomocysteine (SAH) for 1 hour on ice. Nucleosomes containing H3K14ub were purchased from EpiCypher (SKU: 16-0398). The complex was injected onto a Superose 6 Increase 3.2/300 column (Cytiva) using an ÄKTA Micro FPLC system. Fractions containing the complex at 430 nM concentration were crosslinked with 0.1% (v/v) glutaraldehyde for 10 minutes on ice, followed by quenching with 2.4 mM aspartate and 2 mM lysine for 10 minutes on ice. Sample was dialyzed for 3 hours against cryo-EM buffer [20 mM HEPES pH 7.4, 1 mM TCEP pH 8.0, 20 mM NaCl, and 1 mM MgCl_2_].

Quantifoil R2/1 copper grids (200 mesh) were glow-discharged for 30 seconds at 15 mA using a Pelco Easiglow plasma discharge system. A volume of 4 µL of dialyzed sample was applied to each grid and incubated for 8 seconds. Grids were then blotted for 3.5 seconds and vitrified by plunge-freezing in liquid ethane using a Leica GP2 EM system operated at 4 °C and 99% humidity.

Cryo-EM data were acquired on a ThermoFisher Titan Krios transmission electron microscope operating at 300 keV and equipped with a Gatan K3 direct electron detector and BioQuantum energy filter. Data collection was automated using SerialEM v4.0.5. Images were recorded at a calibrated pixel size of 0.83 Å with a defocus range of 0.9–2.1 µm. Each movie consisted of 54 frames with a total dose of 55.1 e^−^/Å^2^ and a nominal spherical aberration of 2.7 mm. A total of 13,149 micrographs were collected.

All data analysis was performed in CryoSPARC v4.4.1. After initial exposure curation, 12,873 micrographs were retained. Particles were picked using the blob picker, resulting in 1,981,230 particles. Local motion correction was applied using a 280-pixel box size, followed by 2D classification. A subset of 744,125 particles was used to generate an ab initio reconstruction, which served as input for multiple rounds of heterogeneous refinement to improve particle homogeneity and resolution. The maps showed significant heterogeneity for the visible extranucleosomal density and secondary structure of the extranucleosomal density could not be resolved, indicating high levels of flexibility. The gray-colored map showed best agreement with the HADDOCK derived model. FSC curves, angular distribution plots, and local resolutions were derived from CryoSPARC.

### Silencing assays

Silencing assays were performed using serial dilution spot tests. *S. pombe* cells were grown overnight in YES medium at 30 °C. Cultures were adjusted to OD_600_ = 1, and 1 mL of cells was harvested, washed with 1 mL sterile water, and resuspended in 250 µL of sterile water. A 3 µL aliquot of the undiluted suspension and 10-fold serial dilutions were spotted onto YE plates and incubated at 32 °C for 3–5 days. For *ade6*^+^ silencing assays, plates were subsequently incubated at 4 °C for an additional 2 days to enhance red pigmentation associated with *ade6*^+^ repression.

### Chromatin Immunoprecipitation (ChIP) and quantitative PCR (qPCR)

ChIP was performed as described previously ^44^ with minor modifications. Briefly, 45 mL of *S. pombe* cells (OD_600_ = 2) were crosslinked with 1% formaldehyde for 15 min at room temperature, quenched with 0.1 M glycine, washed twice with ice-cold TBS, and stored at –80 °C. Cells were lysed in ice-cold lysis buffer [50 mM HEPES-KOH pH 7.5, 140 mM NaCl, 1 mM EDTA, 1% Triton X-100, 0.1% SDS, 0.1% sodium deoxycholate, protease inhibitors, and 1 mM PMSF] using 0.5 mm glass beads in a MagNA Lyser. Lysates were sheared using a Covaris E220 evolution sonicator and clarified by centrifugation at 16000g for 20 minutes at 4°C. 5% of lysate was set aside as input. The remainder was incubated with 2 µg of antibody (H3K9me2, Abcam ab1220; FLAG, Sigma F1804) pre-bound to Protein A or G Dynabeads (Thermo Fisher) for 3 h at 4 °C. Beads were washed three times with lysis buffer and once with TE. Immunoprecipitated chromatin was eluted in two steps using elution buffer [50 mM Tris pH 8.0, 10 mM EDTA, 1% SDS] and [1× TE, 0.67% SDS], followed by the addition of 120 ug RNaseA per mL and. overnight incubation at 65 °C to reverse crosslinks. Samples were treated with proteinase K (360 ug/mL), followed by LiCl/TE extraction, phenol–chloroform purification, and ethanol precipitation. DNA was resuspended in 10 mM Tris-HCl (pH 8.0).

For H3K9me3 ChIP, antibody (Diagenode C15500003) was conjugated to Dynabeads M280 Streptavidin (Invitrogen) per the manufacturer’s instructions. qPCR primers used in this study are listed in Table S4. qPCR was performed using SYBR Green and QuantStudio 7 Flex (Applied Biosystems). The ΔCt method was used to calculate percent input. Data represent the mean of three biological replicates with standard deviations.

### ChIP-seq library preparation and analysis

ChIP-seq libraries were prepared following the protocol described in ^44^, with minor modifications. Briefly, end repair and 5′ phosphorylation were performed on ChIP and input DNA, followed by purification using NucleoSpin columns (Takara). These columns were also used for clean-up between all subsequent enzymatic steps. A-overhangs were added, adapters were ligated, and libraries were PCR-amplified. DNA was purified after each step. Final libraries were quantified using a Qubit fluorometer (Thermo Fisher) and quality assessed with an Agilent TapeStation. Sequencing was performed on an Illumina HiSeq platform.

Sequencing data quality was assessed using *FastQC*. Barcodes were demultiplexed using *FASTX Toolkit*, and adapter sequences were removed with *Cutadapt*. Reads were aligned to the reference genome using *Bowtie2*. Aligned data were converted into BigWig format for visualization in the IGV genome browser.

---

**Figure S1.**
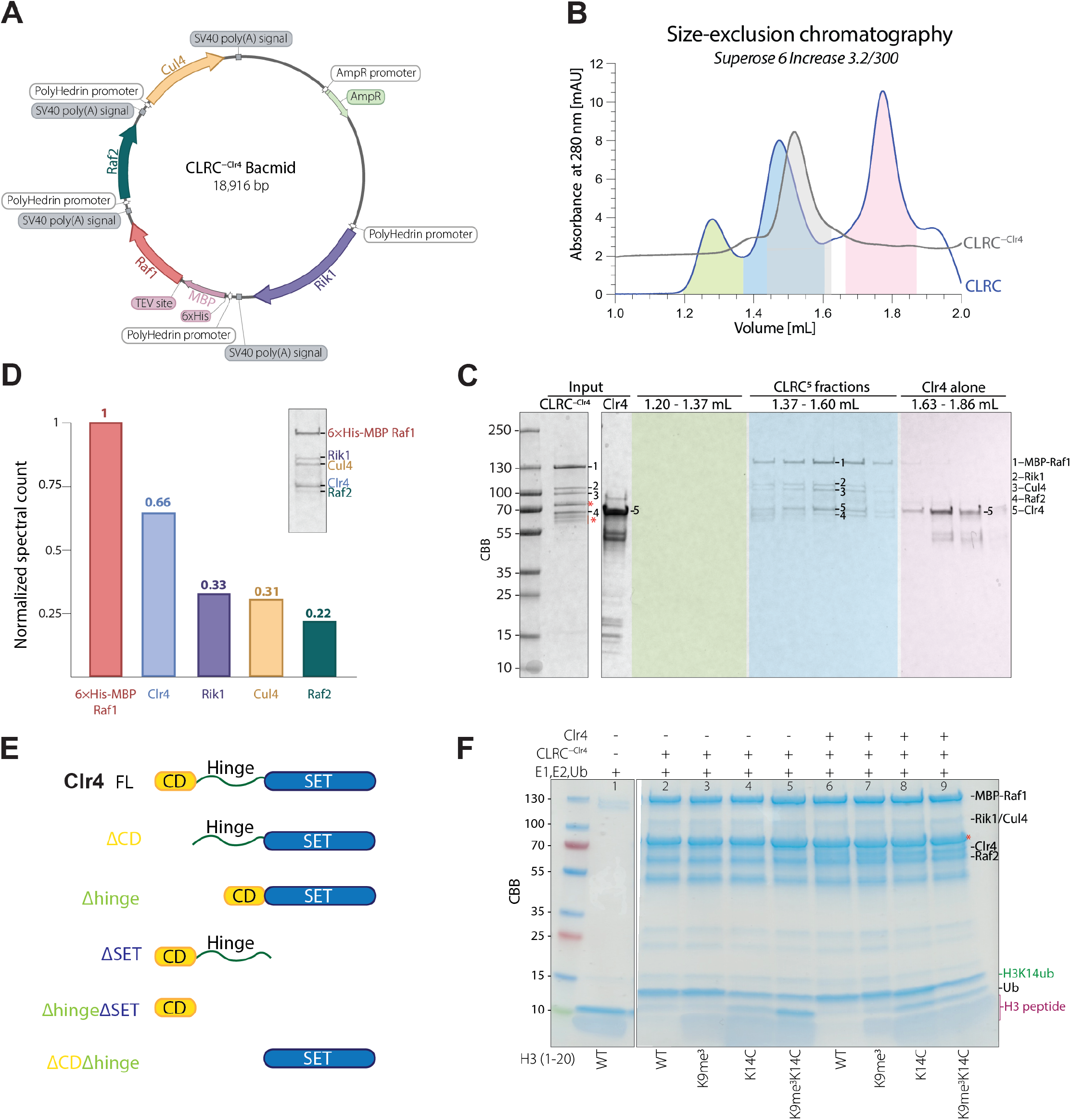
Purification of the CLRC complex. **(A)** Map of Recombinant Bacmid Vector Encoding CLRC^-Clr4^. Diagram of the recombinant bacmid vector used for insect cell expression, encoding all CLRC components except Clr4. Key vector features and regulatory elements are indicated. **(B)** SEC chromatogram of CLRC complexes using Superose 6 Increase 3.2/300 column (Cytiva). Absorbance at 280 nm (mAU) is shown. The grey line indicated the elution profile of CLRC complex lacking Clr4, with the peak fractions highlighted. The blue line indicated the elution profile of the full CLRC complex with peak fractions highlighted in blue and fractions containing excess unbound Clr4 highlighted in pink. **(C)** SDS-PAGE analysis of SEC fractions from panel B. Coomassie-stained SDS–PAGE gel showing input (CLRC^-Clr4^ and Clr4) and eluted fractions from size-exclusion chromatography. Fractions corresponding to the blue peak (1.45–1.55 mL) contain all 5 CLRC complex subunits, while later fractions highlighted in pink (1.65–1.85 mL) contain excess unbound Clr4. *, MBP-Raf1 degradation products. **(D)** Mass spectrometry analysis of the reconstituted CLRC complex. Quantitative LC-MS/MS analysis of the reconstituted CLRC complex showing normalized spectral counts relative to MBP–Raf1 (set to 1). The calculated molar ratios of 0.66 for Clr4, 0.33 for Rik1, 0.31 for Cul4, and 0.22 for Raf2 are consistent with SDS–PAGE band intensities. **(E)** Schematic representation of Clr4 truncation constructs used in this study. Domain architecture of Clr4 and its truncation variants, with the chromodomain shown in yellow, the hinge region in green, and the SET domain in blue. **(F)** SDS-PAGE analysis of in vitro H3 ubiquitination by reconstituted CLRC complex. Ubiquitination reactions were performed using recombinant human E1, E2, HA–ubiquitin (Ub), and the N-terminal peptide of histone H3 (residues 1– 20), in the presence of reconstituted CLRC complexes with or without Clr4. Both CLRC and CLRC^-Clr4^ efficiently ubiquitinated the H3 peptide, whereas E1 and E2 alone showed no activity. *, MBP-Raf1 degradation products; HO, histone octamer.

**Figure S2.**
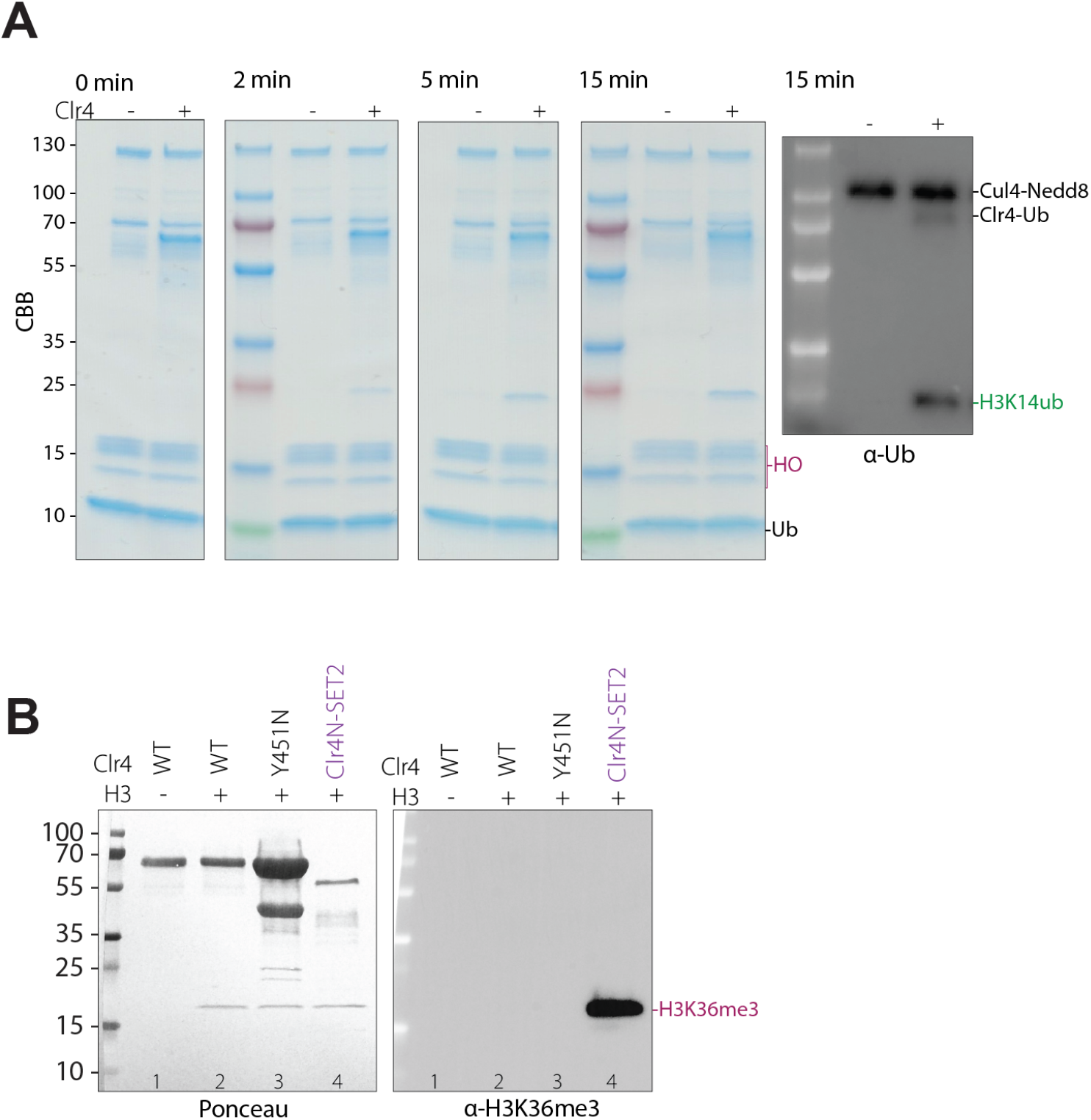
The kinetics of H3K14 ubiquitination and its Clr4-dependence. **(A)** SDS–PAGE analysis of time-course ubiquitination of nucleosome by reconstituted CLRC ± Clr4 complex. Reactions were sampled at the indicated time points to monitor the progression of nucleosome ubiquitination. HO, histone octamer. **(B)** Clr4N-Set2 fusion protein retains H3K36 methylation specificity. Immunoblotting with anti-H3K36me3 antibody demonstrating that the Clr4N-Set2 fusion methylated H3K36.

**Figure S3.**
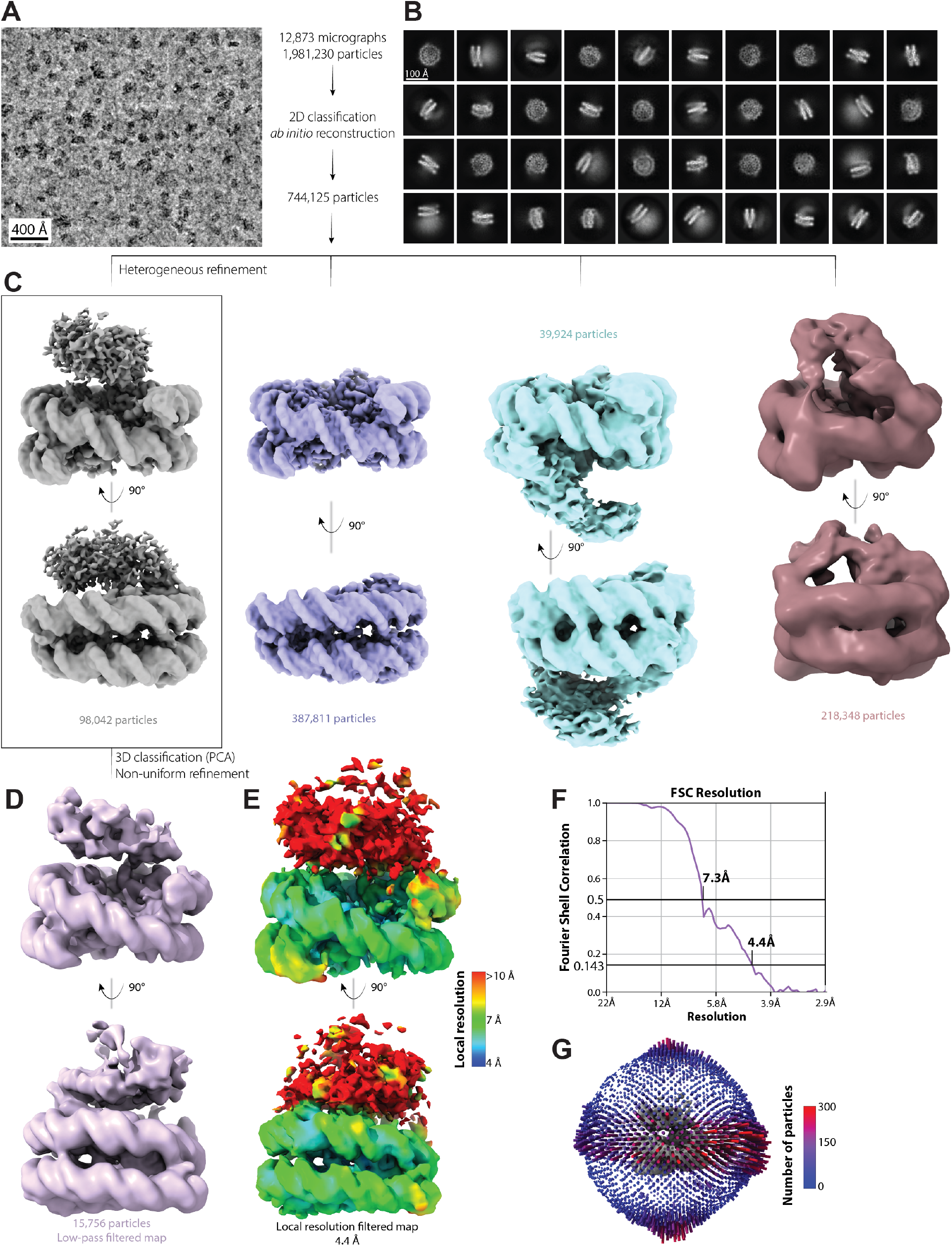
CryoEM data acquisition and processing workflow. Cryo-EM data were collected on a Titan Krios (300 keV) with a Gatan K3 detector. **(A)** Representative low-pass filtered micrograph (upper left; scale bar 400 Å) and classification tree show monodisperse particles. Of 13,149 micrographs, 12,873 were retained after curation. Image processing in CryoSPARC v4.4.1 included blob picking (1,981,230 particles), local motion correction (280-pixel box), and 2D classification. **(B)** Representative 2D class averages (scale bar 100 Å). **(C)** A total of 744,125 particles were used for ab initio reconstruction and multiple rounds of heterogeneous refinement. The map (gray) used for further processing is highlighted by a rectangle. The gray map showed best agreement with the HADDOCK derived model. **(D)** The final low-pass filtered map used for model building, comprising 15,756 particles, was selected from 3D classification based on principal component analysis (PCA). **(E)**The final map was further processed using non-uniform refinement with local resolution filtering. The overall resolution, estimated by FSC, was 4.4 Å. Local resolution analysis (colored from red [low] to blue [high]) revealed that the additional density was of substantially lower resolution, with the global resolution value largely reflecting the high-resolution reconstruction of the nucleosome core. **(F)** FSC curve of the final map from panel E. **(G)** Angular particle distribution plot of the final map from panel E.

**Figure S4.**
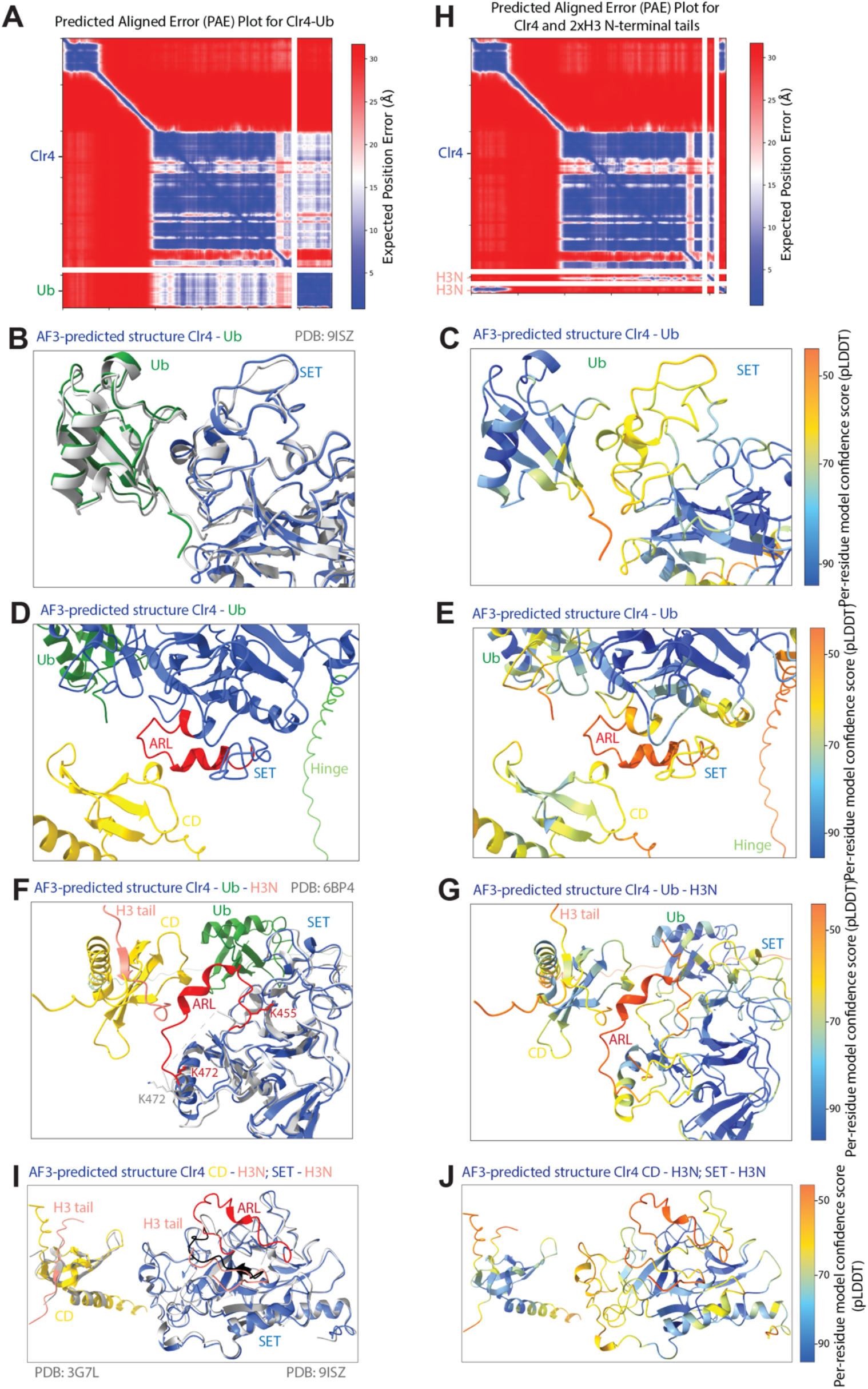
AlphaFold3-based structural predictions of Clr4 and its interactions with ubiquitin and histone H3 tails. **(A)** Predicted Aligned Error (PAE) plots for AlphaFold3 (AF3) model of Clr4-ubiquitin. PAE plot show the predicted alignment error between residue pairs for the Clr4ubiquitin complex. Low predicted error (blue) indicates high confidence in residue placement; high error (white/red) reflects flexibility or lack of interaction. **(B)** Overlay of the AF3-predicted Clr4–ubiquitin complex with the crystal structure of Clr4-SET bound to an H3K14ub peptide. The predicted interface between the Clr4 SET domain (blue) and ubiquitin (green) closely matches the crystal structure (PDB 9ISZ, grey). The corresponding PAE plot is shown in panel A. **(C)** Per-residue model confidence (pLDDT) for the AF3 prediction in panel B. High-confidence regions are indicated in blue; lower-confidence regions are shown in green to red. **(D)** AF3-predicted intramolecular interaction between Clr4’s autoregulatory loop (ARL, red) and chromodomain (CD, yellow). **(E)** pLDDT scores for the predicted structure in panel D. **(F, G)** Overlay of the predicted Clr4 SET domain structure with the crystal structure of open automethylated Clr4 SET domain. The AF3-predicted structure (blue) aligns closely with the SET domain in its open conformation (PDB 6BP4, grey). pLDDT confidence scores are shown in panel G and the PAE plot in panel A. **(H)** Predicted Aligned Error (PAE) plots for AlphaFold3 (AF3) models of Clr4-H3 tail. **(I, J)** Overlay of the AF3-predicted interactions between Clr4 and H3 tails with experimental structures. AF3-predicted binding of the Clr4 CD (yellow) and SET domain (blue) to the H3 N-terminal tail (salmon) is in close agreement with respective crystal structures of CD-H3 (PDB 3G7L, grey) and SET–H3 (PDB 9ISZ, grey) (I). pLDDT confidence scores are shown in panel J and the PAE plot in panel H.

**Figure S5.**
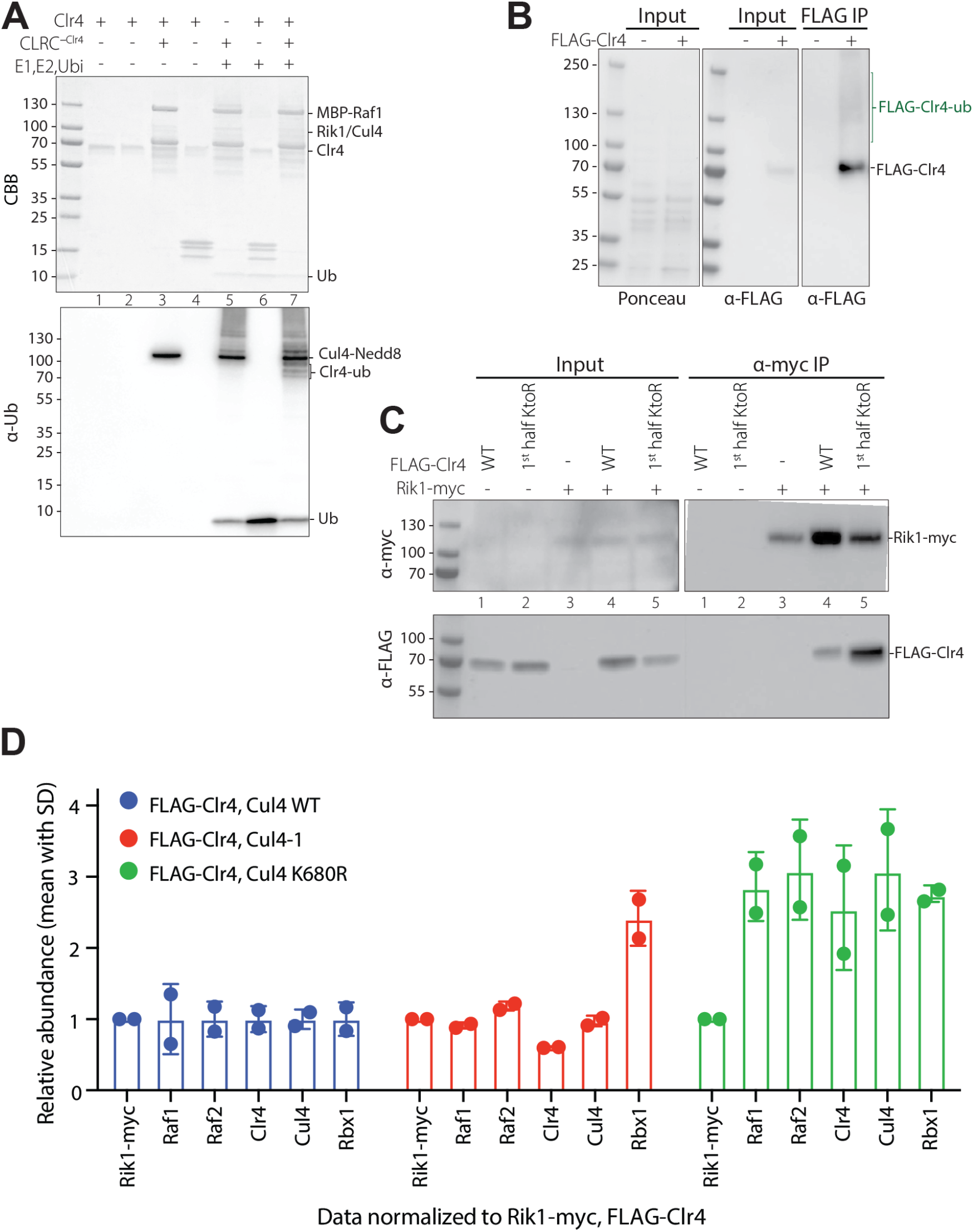
CLRC mediated Clr4 ubiquitination and release of Clr4 from the CLRC complex. **(A)** CLRC-dependent ubiquitination of Clr4 in vitro. Anti-ubiquitin western blot of in vitro reactions using purified CLRC complexes. Mono- and di-ubiquitinated Clr4 species (lane 7) appeared only in the presence of the full CLRC complex, but not with E1/E2 enzymes or CLRC lacking Clr4 (lanes 5–6), indicating CLRC-dependent ubiquitination. The anti-ubiquitin antibody also cross-reacts with Nedd8, resulting in a detectable Cul4-Nedd8 band. (**B**) Detection of polyubiquitinated Clr4 species in vivo. Western blot analysis of FLAG-Clr4 purified from *S. pombe* in the presence of the proteasome inhibitor MG-132 and deubiquitinase inhibitor NEM revealed slower-migrating bands, consistent with polyubiquitinated forms of Clr4. **(C)** Loss of Clr4 ubiquitination enhances interaction with CLRC. Western blot analysis of co-immunoprecipitation experiments in *S. pombe* strains expressing myc-tagged Rik1 and FLAG-Clr4 variants. Immunoprecipitation with anti-myc beads revealed that Clr4 1^st^ half KtoR mutants, which lack ubiquitination in the chromodomain and hinge regions, exhibited increased association with the CLRC complex compared to wild-type Clr4. **(D)** Quantitative proteomic analysis of CLRC complex assembly in *cul4* mutant backgrounds. TMT-based mass spectrometry of FLAG-Clr4 immunoprecipitates from *S. pombe* cells expressing wild-type *cul4, cul4-1*, or *cul4-K680R*. In *cul4-K680R* mutant cells, FLAG-Clr4 showed ~3-fold increased association with other CLRC subunits, indicating impaired disassembly. By contrast, the *cul4-1* hypomorphic allele reduced Clr4 association with Rik1 and overall CLRC assembly, consistent with decreased ubiquitination activity.

**Figure S6.**
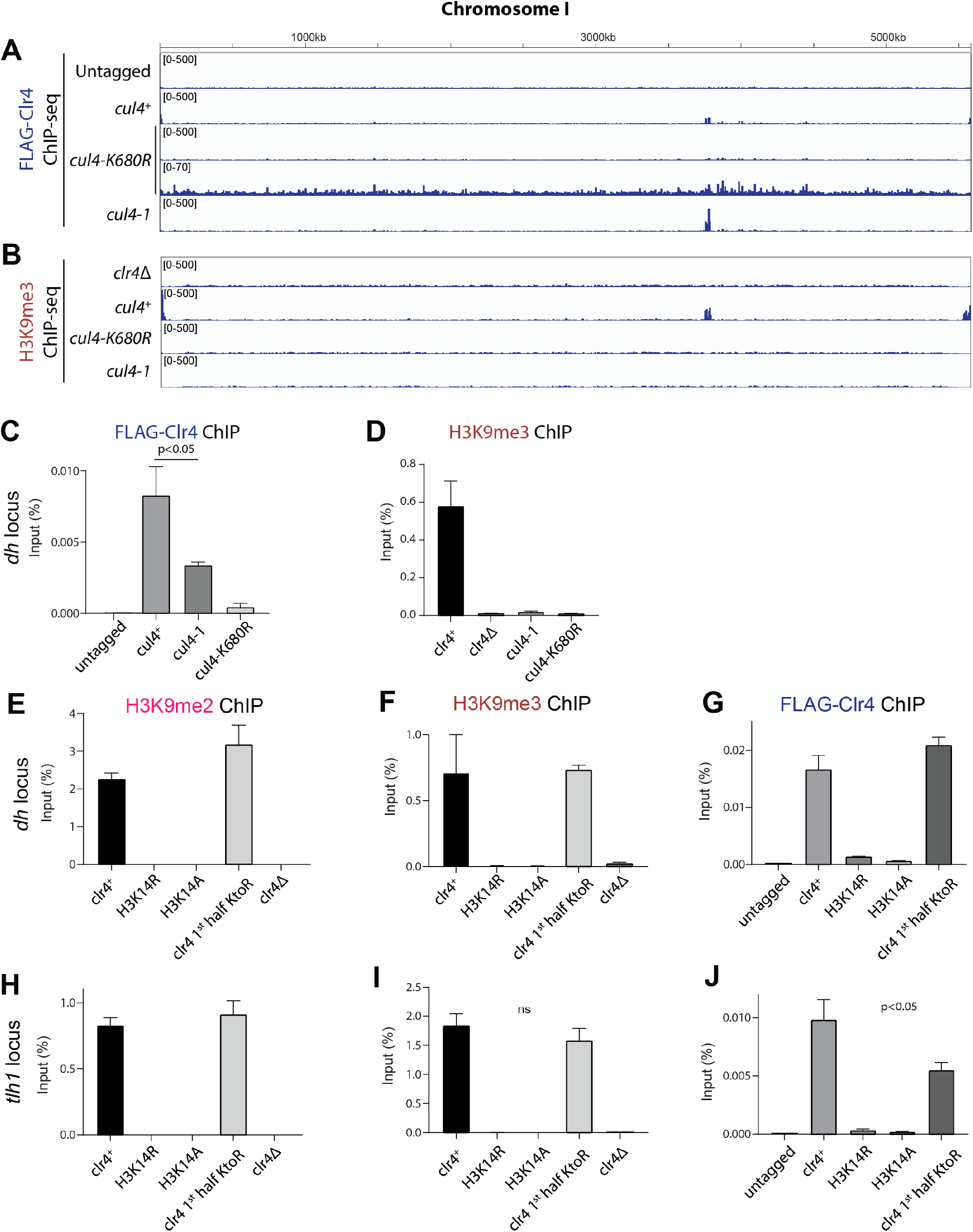
Role of Cul4-mediated ubiquitination in Clr4 localization and H3K9 methylation. Related to Figure 6. (**A, B**) Genome browser views of ChIP-seq data showing the localization of FLAG-Clr4 (A) and H3K9me3 (B) along the *S. pombe* chromosome I in wild-type (*cul4*^*+*^), *cul4-K680R*, and *cul4-1* cells. **(C-J)** ChIP-qPCR analysis of FLAG-Clr4 and H3K9me3 in wild-type, *cul4-K680R, cul4-1, H3K14R, H3K14A, 1*^*st*^ *half-KtoR* (clr4 with 1^st^ half lysines substituted with arginine, see Figure 5B), and *clr4*Δ cells at the indicated loci. Bars show mean percentage input and error bars show standard deviations of 3 biological replicates.

**Figure S7.**
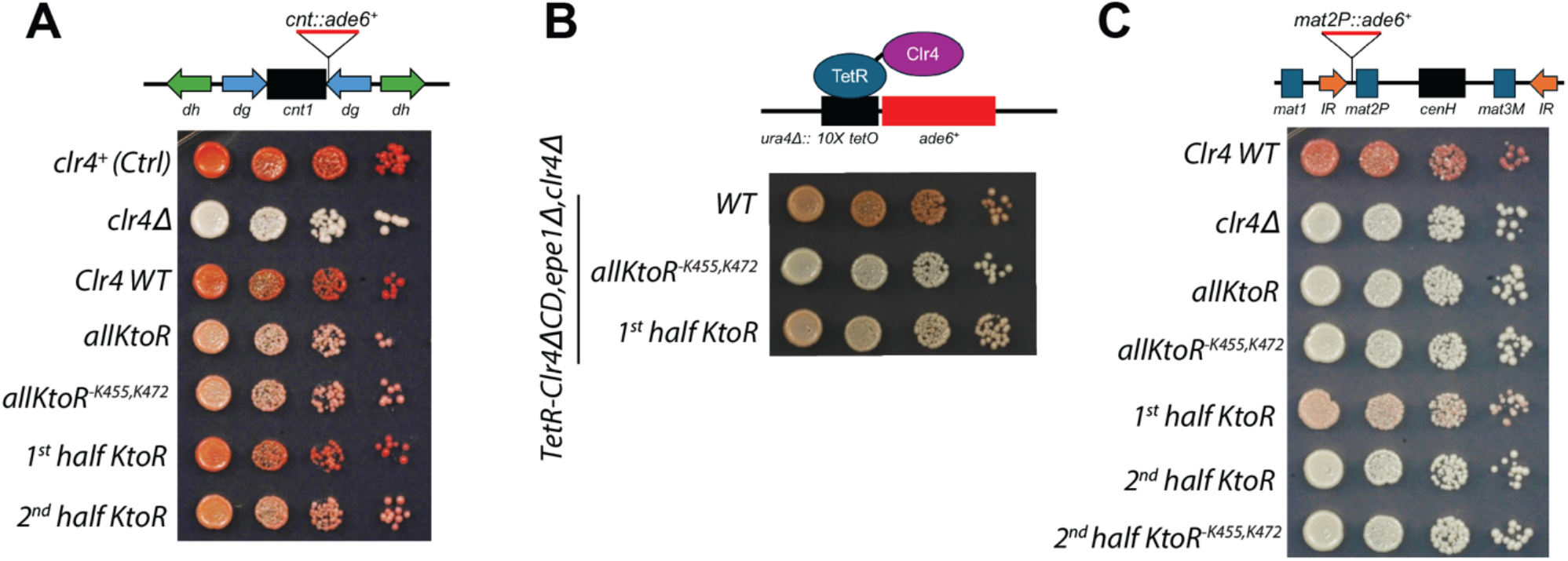
Role of Cul4-mediated ubiquitination in silencing. (**A**) Silencing assays for a centromeric *ade6*^+^ reporter (*otr1R::ade6*^*+*^). Ten-fold serial dilutions of cells were plated on low adenine YE plates in which silencing of *ade6*^+^ leads to formation of red/pink colonies. (**B**) Silencing assays for the *ade6*^+^ reporter inserted at the mating type locus (*mat2P::ade6*^+^). Cells were plated as in panel A. **(C)** Silencing assays for the *ura4Δ::10XtetO-ade6*^+^ reporter cells in which the recruitment of TetR-Clr4-ΔCD initiator leads to formation of heterochromatin and *ade6*^+^ silencing. Wild-type (WT) refers to TetR-Clr4-ΔCD without any amino acid substitutions.

**Figure S8.**
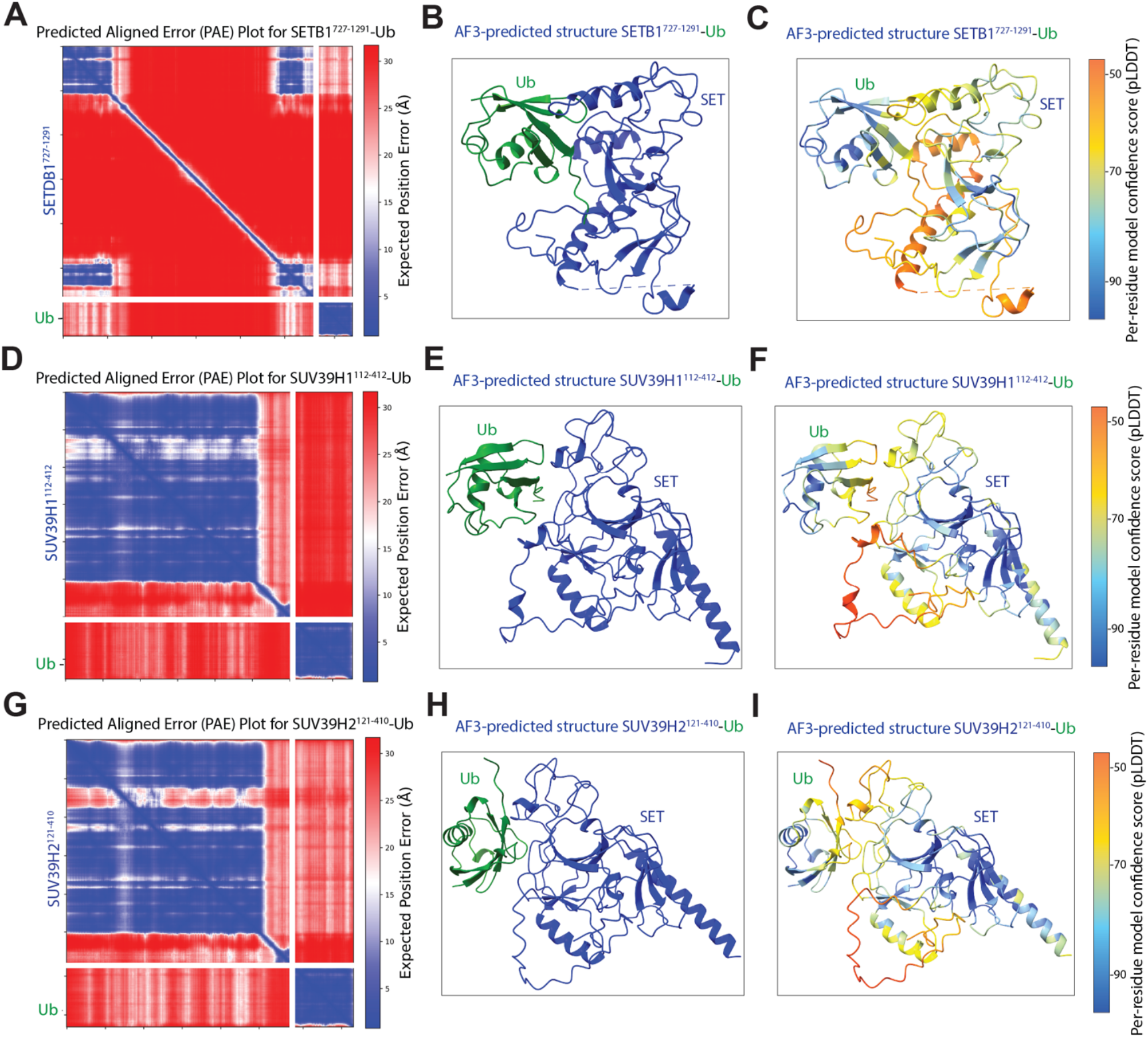
AlphaFold3 structural predictions of the interactions of SUV39 subfamily of SET-domain-containing histone methyltransferases with ubiquitin. **(A, D, G)** Predicted Aligned Error (PAE) plots for AlphaFold3 (AF3) models of the indicated SET domains and ubiquitin (Ub). PAE plots show the predicted alignment error between residue pairs for the SETDB1^727-1291^-ubiquitin complex (A), SUV39H1^112-412^-ubiquitin complex (D), SUV39H2^121-410^-ubiquitin complex (G). Low predicted error (blue) indicates high confidence in residue placement; high error (white/red) reflects flexibility or lack of predicted interaction. **(B, E, H)** AF3-predicted interactions between SUV39 subfamily of SET-domain-containing histone methyltransferases and ubiquitin. The SET domains are highlighted in blue, ubiquitin in green. **(C, F, I)** Per-residue model confidence (pLDDT) for the AF3 predictions in panels B, E, H. High-confidence regions are indicated in blue; lower-confidence regions are shown in green to red.

**Table S1.** Summary of inter- and intra-Clr4 cross-links identified by BS3 XL-MS. All cross-links detected with Clr4 using the amine-reactive cross-linker BS3, which links lysine side chains or N-terminal α-amino groups. The BS3 spacer arm, together with lysine side chains, allows for cross-links up to approximately 30 Å in Cα-Cα distance. To assess structural compatibility, each cross-link was mapped onto the structural model and color-coded by the measured Cα-Cα distance: green: distances ≤ 30 Å, considered compatible with cross-linking constraints; pink: distances between 31-34 Å, likely reflecting flexible regions; dark red: distances > 35 Å, exceeding the cross-linker limit and indicating possible violations or conformational variability. Structural domains and regions are annotated as follows: chromodomain (CD) in yellow, hinge region in light green, SET domain in blue, ARL in red, ubiquitin (Ub) in dark green, and H3 N-terminal tail in purple. All identified crosslinks, including those within the nucleosome, are reported in Table S5.

| Protein #1 | Amino acid #1 | Residue number #1 | Protein #2 | Amino acid #2 | Residue number #2 | Crosslinked regions | Distance |
| --- | --- | --- | --- | --- | --- | --- | --- |
| Clr4 | GLY | 1 | Clr4 | LYS | 17 | CD-CD | 17 Å |
| Clr4 | GLY | 1 | Clr4 | LYS | 25 | CD-CD | 18 Å |
| Clr4 | GLY | 1 | Clr4 | LYS | 94 | CD-hinge | 26 Å |
| Clr4 | GLY | 1 | Clr4 | LYS | 109 | CD-hinge | 29 Å |
| Clr4 | GLY | 1 | Clr4 | LYS | 114 | CD-hinge | 27 Å |
| Clr4 | GLY | 1 | Clr4 | LYS | 122 | CD-hinge | 20 Å |
| Clr4 | GLY | 1 | Clr4 | LYS | 187 | CD-hinge | 39 Å |
| Clr4 | GLY | 1 | Clr4 | LYS | 472 | CD-ARL | 24 Å |
| Clr4 | LYS | 17 | Clr4 | LYS | 25 | CD-CD | 6 Å |
| Clr4 | LYS | 17 | Clr4 | LYS | 472 | CD-ARL | 24 Å |
| Clr4 | LYS | 109 | Clr4 | LYS | 114 | Hinge-hinge | 11 Å |
| Clr4 | LYS | 122 | Clr4 | LYS | 160 | Hinge-hinge | 18 Å |
| Clr4 | LYS | 151 | Clr4 | LYS | 160 | Hinge-hinge | 16 Å |
| Clr4 | LYS | 151 | Clr4 | LYS | 165 | Hinge-hinge | 25 Å |
| Clr4 | LYS | 154 | Clr4 | LYS | 165 | Hinge-hinge | 18 Å |
| Clr4 | LYS | 160 | Clr4 | LYS | 174 | Hinge-hinge | 25 Å |
| Clr4 | LYS | 165 | Clr4 | LYS | 174 | Hinge-hinge | 14 Å |
| Clr4 | LYS | 165 | Clr4 | LYS | 472 | Hinge-ARL | 23 Å |
| Clr4 | LYS | 174 | Clr4 | LYS | 187 | Hinge-hinge | 6 Å |
| Clr4 | LYS | 174 | Clr4 | LYS | 472 | Hinge-ARL | 20 Å |
| Clr4 | LYS | 180 | Clr4 | LYS | 193 | Hinge-SET | 24 Å |
| Clr4 | LYS | 187 | Clr4 | LYS | 193 | Hinge-SET | 17 Å |
| Clr4 | LYS | 187 | Clr4 | LYS | 205 | Hinge-SET | 21 Å |
| Clr4 | LYS | 187 | Clr4 | LYS | 472 | Hinge-ARL | 25 Å |
| Clr4 | LYS | 193 | Clr4 | LYS | 211 | SET-SET | 26 Å |
| Clr4 | LYS | 334 | Clr4 | LYS | 338 | SET-SET | 8 Å |
| Clr4 | LYS | 372 | Clr4 | LYS | 472 | SET-ARL | 20 Å |
| Clr4 | LYS | 464 | Clr4 | LYS | 472 | ARL-ARL | 17 Å |
| Clr4 | LYS | 464 | Clr4 | LYS | 478 | ARL-ARL | 16 Å |
| Clr4 | LYS | 472 | Clr4 | LYS | 478 | ARL-ARL | 19 Å |
| Clr4 | GLY | 1 | H2A | LYS | 95 | CD-H2A | 25 Å |
| Clr4 | LYS | 154 | H2A | LYS | 9 | CD-H2A | 30 Å |
| Clr4 | LYS | 154 | H2A | LYS | 36 | CD-H2A | 27 Å |
| Clr4 | LYS | 165 | H2A | LYS | 36 | CD-H2A | 32 Å |
| Clr4 | LYS | 187 | H2A | LYS | 36 | CD-H2A | 34 Å |
| Clr4 | LYS | 472 | H2A | LYS | 36 | ARL-H2A | 36 Å |
| Clr4 | GLY | 1 | H2B | LYS | 31 | CD-H2B | 44 Å |
| Clr4 | GLY | 1 | H2B | LYS | 105 | CD-H2B | 18 Å |
| Clr4 | GLY | 1 | H2B | LYS | 113 | CD-H2B | 27 Å |
| Clr4 | GLY | 1 | H2B | LYS | 117 | CD-H2B | 32 Å |
| Clr4 | LYS | 17 | H2B | LYS | 105 | CD-H2B | 28 Å |
| Clr4 | LYS | 17 | H2B | LYS | 117 | CD-H2B | 36 Å |
| Clr4 | LYS | 25 | H2B | LYS | 105 | CD-H2B | 31 Å |
| Clr4 | LYS | 109 | H2B | LYS | 105 | Hinge-H2B | 32 Å |
| Clr4 | LYS | 113 | H2B | LYS | 117 | Hinge -H2B | 25 Å |
| Clr4 | LYS | 114 | H2B | LYS | 8 | Hinge -H2B | 56 Å |
| Clr4 | LYS | 122 | H2B | LYS | 113 | Hinge -H2B | 24 Å |
| Clr4 | LYS | 122 | H2B | LYS | 117 | Hinge -H2B | 26 Å |
| Clr4 | LYS | 154 | H2B | LYS | 117 | Hinge -H2B | 15 Å |
| Clr4 | LYS | 154 | H2B | LYS | 122 | Hinge -H2B | 20 Å |
| Clr4 | LYS | 174 | H2B | ALA | 1 | Hinge -H2B | 25 Å |
| Clr4 | LYS | 180 | H2B | ALA | 1 | Hinge -H2B | 13 Å |
| Clr4 | LYS | 187 | H2B | ALA | 1 | Hinge -H2B | 23 Å |
| Clr4 | LYS | 464 | H2B | ALA | 1 | ARL-H2B | 45 Å |
| Clr4 | GLY | 1 | H3 | LYS | 4 | CD-H3 | 30 Å |
| Clr4 | GLY | 1 | H3 | LYS | 18 | CD-H3 | 10 Å |
| Clr4 | GLY | 1 | H3 | LYS | 23 | CD-H3 | 11 Å |
| Clr4 | GLY | 1 | H3 | LYS | 27 | CD-H3 | 21 Å |
| Clr4 | GLY | 1 | H3 | LYS | 56 | CD-H3 | 42 Å |
| Clr4 | GLY | 1 | H3 | LYS | 79 | CD-H3 | 15 Å |
| Clr4 | LYS | 17 | H3 | LYS | 4 | CD-H3 | 24 Å |
| Clr4 | LYS | 17 | H3 | LYS | 27 | CD-H3 | 31 Å |
| Clr4 | LYS | 25 | H3 | LYS | 27 | CD-H3 | 29 Å |
| Clr4 | LYS | 94 | H3 | LYS | 4 | Hinge-H3 | 19 Å |
| Clr4 | LYS | 127 | H3 | LYS | 27 | Hinge-H3 | 31 Å |
| Clr4 | LYS | 154 | H3 | LYS | 23 | Hinge-H3 | 32 Å |
| Clr4 | LYS | 154 | H3 | LYS | 122 | Hinge-H3 | 45 Å |
| Clr4 | LYS | 211 | H3 | LYS | 4 | SET-H3 | 37 Å |
| Clr4 | LYS | 338 | H3 | LYS | 4 | SET-H3 | 27 Å |
| Clr4 | LYS | 372 | H3 | LYS | 4 | SET-H3 | 4 Å |
| Clr4 | LYS | 464 | H3 | LYS | 4 | ARL-H3 | 24 Å |
| Clr4 | LYS | 464 | H3 | LYS | 18 | ARL-H3 | 17 Å |
| Clr4 | LYS | 464 | H3 | LYS | 23 | ARL-H3 | 16 Å |
| Clr4 | LYS | 464 | H3 | LYS | 27 | ARL-H3 | 21 Å |
| Clr4 | LYS | 472 | H3 | LYS | 4 | ARL-H3 | 18 Å |
| Clr4 | LYS | 472 | H3 | LYS | 27 | ARL-H3 | 38 Å |
| Clr4 | LYS | 472 | H3 | LYS | 122 | ARL-H3 | 32 Å |
| Clr4 | GLY | 1 | H4 | LYS | 20 | CD-H4 | 15 Å |
| Clr4 | GLY | 1 | H4 | LYS | 31 | CD-H4 | 30 Å |
| Clr4 | GLY | 1 | H4 | LYS | 91 | CD-H4 | 31 Å |
| **Clr4** | **LYS** | **17** | **H4** | **LYS** | **20** | **CD-H4** | **31 Å** |
| **Clr4** | **GLY** | **1** | **Ub** | **LYS** | **6** | **CD-Ub** | **8 Å** |
| **Clr4** | **GLY** | **1** | **Ub** | **LYS** | **48** | **CD-Ub** | **19 Å** |
| **Clr4** | **LYS** | **17** | **Ub** | **LYS** | **48** | **CD-Ub** | **12 Å** |
| **Clr4** | **LYS** | **154** | **Ub** | **LYS** | **48** | **Hinge-Ub** | **31 Å** |
| **Clr4** | **LYS** | **472** | **Ub** | **LYS** | **6** | **ARL-Ub** | **24 Å** |

**Table S2.** List of *S. pombe* strains used in this study.

| Strain # | Genotype | Source |
| --- | --- | --- |
| **SPY3** | *mat1Msmto leu1-32 his2 L(Bgl II)::ade6+ ade6 DN/N* | S.I. Grewal |
| **SPY7283** | *SPY3 hphR-5'(1kb)-3xFlag-clr4+ #1* | Lab stock |
| **SPY7305** | *SPY3 clr4Δ::kanR #1* | Lab stock |
| **SPY11057** | *SPY3 clr4 all KR #1* | this study |
| **SPY11058** | *SPY3 clr4 all KR except K455/K472 #1* | this study |
| **SPY11059** | *SPY3 clr4 1st half KR #2* | this study |
| **SPY11071** | *SPY3 hphR-Flag-clr4 2nd half KR #7* | this study |
| **SPY10562** | *SPY3 hphR-FLAG-Clr4-2ndhalf-KtoR_except K455/K472* | this study |
| **SPY5071** | *h- leu1-32 ade6-M210 ura4Δ::10XTetO-ade6 #1* | Lab stock |
| **SPY5086** | *h- leu1-32 ade6+-M210 ura4Δ::10XTetO- ade6+ clr4Δ::natMX6-Pclr4-TetR-2XFLAG-clr4-I epe1Δ::kanMX6* | Lab stock |
| **SPY10655** | *h- leu1-32 ade6-M210 ura4Δ::10XTetO-ade6 clr4Δ::nat-clr4p-TetR-Clr4-allKtoR-except K455,K472-ΔCD4* | this study |
| **SPY10657** | *h- leu1-32 ade6-M210 ura4Δ::10XTetO-ade6 clr4Δ::nat-clr4p-TetR-Clr4-1st half-KtoR-ΔCD* | this study |
| **SPY138** | *h+ leu1-32 ade6-M210 ura4-D18 otr1R(SphI)::ade6+* | K. Ekwall |
| **SPY8328** | *SPY138 clr4Δ::ura4-KAN #1* | Lab stock |
| **SPY5233** | *SPY138 Kan-1kb5'UTR-3xFlag-clr4 #1* | Lab stock |
| **SPY11061** | *SPY138 hphR-Flag-clr4 all KR #21* | this study |
| **SPY11063** | *SPY138 hphR-Flag-clr4 all KR except K455/K472* | this study |
| **SPY11065** | *SPY138 hphR-Flag-clr4 1st half KR* | this study |
| **SPY11067** | *SPY138 hphR-Flag-clr4 2nd half KR* | this study |
| **SPY11277** | *h+ leu1-32 ade6-M210 ura4DS/E otr1R(SphI)::ura4+ oriA FLAG-clr4 Rik1-9myc cul4-GFP* | this study |
| **SPY11278** | *h+ leu1-32 ade6-210 ura5-D18 otr1R(SphI)::ade6+ 3xflag-clr4, Rik1-9myc::kan, cul4680R::Nat* | this study |
| **SPY1702** | *6079 h3.2-K14R h3.1/h4.1 ::his3+ h3.3/h4.3 ::arg3+ ade6-210 otr1R(SphI):ade6+* | R. Allshire |
| **SPY1700** | *6076 h3.2-K14A h3.1/h4.1 ::his3+ h3.3/h4.3 ::arg3+ ade6-210 otr1R(SphI):ade6+* | R. Allshire |
| **SPY10829** | *h+ h3.2-K14R h3.1/h4.1::his3+ h3.3/h4.3::arg3+ Kan-1kb5'UTR-3xFlag-Clr4 ade6-M210 otr1R(SphI):ade6+* | this study |
| **SPY10825** | *h+ h3.2-K14A h3.1/h4.1::his3+ h3.3/h4.3::arg3+ Kan-1kb5'UTR-3xFlag-Clr4 ade6-M210 otr1R(SphI):ade6+* | this study |
| **SPY137** | *h+ leu1-32 ade6-M210 ura4DS/E otr1R(SphI)::ura4+ oriA* | K. Ekwall |
| **SPY815** | *SPY137 clr4Δ::kanR* | Hong et al. |

**Table S3.** List of plasmids used in this study.

| **Plasmid #** | **Description** | **Source** |
| --- | --- | --- |
| **pDM2113** | pGEX-6P-1-Clr4 WT | Lab stock |
| **pDM2142** | pGEX-6P-1-Clr4-Y451N | Lab stock |
| **pDM1000** | pGEX-6P-1-Clr4 ΔCD (aa 70 - 490) | Lab stock |
| **pDM1906** | pGEX-6P-1-Clr4 SET (aa 192-490) | Lab stock |
| **pDM2499** | pGEX-6P-1-Clr4 ΔHinge (Δaa 70-191) 2×GSS | This study |
| **pDM2426** | pGEX-6P-1-Clr4 ΔSET (aa 1-192) | This study |
| **pDM2432** | pGEX-6P-1-Clr4 ΔhingeΔSET (aa 1-69) | This study |
| **pDM2428** | pGEX-6P-1-Clr4N-Set2 | This study |
| **pDM2132** | pGEX-6P-1-Clr4-K455,472R | Lab stock |
| **pDM2433** | pGEX-6P-1-Clr4-W31G | This study |
| **pDM2451** | pGEX-6P-1-Clr4-W31G,Y451N | This study |
| **pDM2458** | pREP1-nmt1-LEU2 6×His-Ub | Lab stock |
| **pDM2455** | 438-A | L. Farnung |
| **pDM2456** | 438-C | L. Farnung |
| **pDM2457** | pbiGBac | L. Farnung |
| **pDM844** | pET3a-H2A, *Xenopus laevis* | Lab stock |
| **pDM845** | pET3a-H2B, *Xenopus laevis* | Lab stock |
| **pDM1363** | pET3a-H3 C110A, *Xenopus laevis* | Lab stock |
| **pDM1365** | pET3a-H3 K9C,C110A, *Xenopus laevis* | Lab stock |
| **pDM2452** | pET3a-H3 K14R,C110A, *Xenopus laevis* | This study |
| **pDM2453** | pET3a-H3 K9M,C110A, *Xenopus laevis* | This study |
| **pDM2454** | pET3a-H3K9M,K14R,C110A, *Xenopus laevis* | This study |
| **pDM847** | pET3a-H4, *Xenopus laevis* | Lab stock |

**Table S4.**
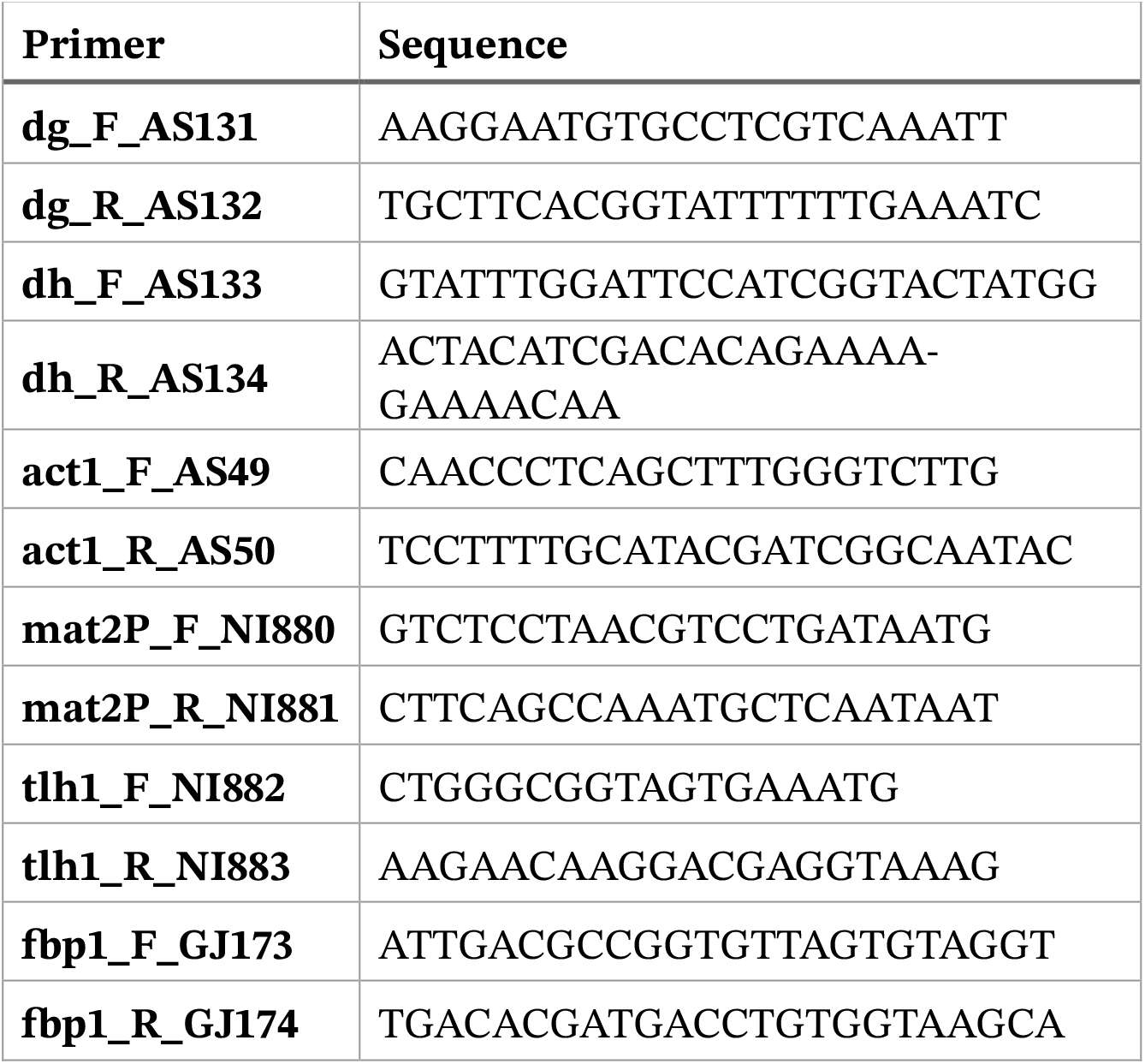
List of qPCR primers used in this study.

